# Loss of NudE-mediated dynein activation at synaptic terminals causes progressive axon length-dependent neurodegeneration

**DOI:** 10.64898/2026.04.12.717987

**Authors:** Zeeshan Mushtaq, Dario Andrea Lasser, Lena Maria Lion, Johannes Ebding, Ben Escribano, Denis Burkhalter, Eliza Moreno, Tanja Maritzen, Raiko Stephan, Jan Pielage

## Abstract

Progressive, axon length-dependent degeneration of nerve terminals is a defining feature of dying-back neuropathies; yet, whether defects in axonal transport initiate or are a consequence of this process remains unresolved. Here, we show that NudE, a scaffold for dynein motor activation, is required for the initiation of retrograde axonal transport at *Drosophila* motoneuron synapses *in vivo*. Loss of NudE impairs dynein activation at the synaptic terminal, severely reducing the proportion of cargo entering retrograde transport and decreasing retrograde motor velocity. Live imaging and temporal analysis establish that this transport initiation defect is the earliest event in a degenerative cascade, followed by progressive microtubule destabilization, impaired synaptic transmission, and structural degeneration in a distal-to-proximal gradient that recapitulates dying-back neuropathies. Structure-function analysis validates the biochemically defined NudE-dynein binding domains *in vivo*, with graded disruption producing correspondingly graded phenotypes. Genetic two-hit experiments uncover a reciprocal dependence between transport initiation and microtubule maintenance: perturbations in either process that are individually tolerated synergistically trigger degeneration when combined, while increasing microtubule levels alone cannot compensate for failed dynein activation. These findings reveal retrograde transport initiation as a critical vulnerability point in motoneurons and provide a mechanistic basis for how defects in dynein activation produce progressive neurodegeneration.

## Introduction

Many neurodegenerative diseases share a common pattern of progression: nerve terminals degenerate first, and the disease advances proximally toward the cell body over time (Moloney et al., 2014; Schaumburg et al., 1974). This dying-back pattern is observed across a broad range of conditions, from inherited peripheral neuropathies such as Charcot-Marie-Tooth disease to motor neuron diseases including amyotrophic lateral sclerosis (Rossor et al., 2012; Sleigh et al., 2019). A defining feature of these diseases is that the longest axons are affected first, suggesting that axon length itself is a vulnerability factor. Microtubules are central to this vulnerability: they serve as the structural backbone of the axon and as tracks for long-range transport of organelles, signaling complexes, and crucial synaptic proteins. Accordingly, mutations in *tubulin* genes (Smith et al., 2014), microtubule-associated proteins (Matamoros and Baas, 2016), and components of the microtubule-based cytoplasmic dynein motor complex, comprising dynein, dynactin and additional regulatory factors all cause progressive neurodegeneration (Farrer et al., 2009; Hafezparast et al., 2003; Puls et al., 2003). Disruption of the dynein-dynactin complex, transporting cargo toward the neuronal cell body, in postnatal motor neurons is sufficient to produce a late-onset degenerative phenotype in mice (LaMonte et al., 2002), directly implicating retrograde axonal transport in neuronal survival. However, whether transport defects are the initiating event in synaptic degeneration or a secondary consequence of progressive cellular dysfunction remains a central open question (Berth and Lloyd, 2023).

To identify microtubule-associated factors whose loss produces length-dependent synaptic degeneration, we performed a genetic screen in *Drosophila* motoneurons. The *Drosophila* larval neuromuscular junction (NMJ) provides an experimentally accessible model in which individual synapses can be analyzed by live imaging, electrophysiology, and high-resolution microscopy, and in which the segmental organization of the body wall creates a natural anterior-posterior gradient of motoneuron axon length. Our screen identified *nudE*, the sole *Drosophila* ortholog of the mammalian dynein regulatory factors Nde1 and Ndel1 (Bradshaw et al., 2013; Wainman et al., 2009). An independent RNAi screen had previously identified *nudE* as required for synaptic maintenance and axonal transport, but the underlying mechanisms were not explored (Valakh et al., 2012). Analysis of nudE in sensory neurons revealed roles in dendrite morphogenesis with comparatively mild axonal phenotypes (Arthur et al., 2015). As the single ortholog of both Nde1 and Ndel1, Drosophila NudE enables an unambiguous genetic analysis that is not confounded by paralog redundancy (Bradshaw et al., 2013; Garrott et al., 2022).

NudE is a regulatory factor of the cytoplasmic dynein motor complex, the sole motor for retrograde axonal transport. In its default state cytoplasmic dynein is autoinhibited through dimerization of its motor domains into a closed conformation termed the phi particle (Zhang et al., 2017). Activation requires the regulatory factor Lis1, which binds dynein and promotes an open conformation competent for assembly with the dynactin complex and a cargo-specific adaptor protein into a processive transport complex (McKenney et al., 2014; Singh et al., 2024). Recent *in vitro* reconstitution studies have revealed that Nde1 acts as a transient scaffold in this process: it tethers Lis1 to autoinhibited dynein through a conserved interaction at the dynein intermediate chain (Okada et al., 2023), enabling the formation of a structural intermediate that is rate-limiting for dynein activation (Yang et al., 2026; Zhao et al., 2023). Lis1 also promotes the assembly of transport complexes containing two dynein dimers, which exhibit higher velocity and force production (Htet et al., 2020). While holding Lis1 and dynein together, Nde1 simultaneously keeps the motor in an inactive state by preventing the interactions required for full complex assembly. Nde1 must be released from the dynein motor domain before the processive dynein-dynactin transport complex can form (Garrott et al., 2023; Zhao et al., 2023). At the synaptic terminal, retrograde transport initiation requires the local assembly of the dynein-dynactin complex on dynamic microtubule plus-ends (Lloyd et al., 2012; Moughamian et al., 2013; Moughamian and Holzbaur, 2012). Dynein and dynactin are delivered to the axon tip by separate anterograde mechanisms and must assemble there (Fellows et al., 2024). Despite the well-characterized biochemistry of this activation cascade, whether NudE/Nde1 is required specifically for transport initiation in neurons *in vivo*, and what the consequences of failed dynein activation are for synaptic integrity, has not been addressed.

Here, we address these questions using the *Drosophila* motoneuron system. We show that NudE is essential for the initiation of retrograde transport at the synaptic terminal and that its loss triggers a progressive degenerative cascade in which transport initiation failure, microtubule destabilization, and synaptic degeneration can be resolved as temporally and genetically separable events. Structure-function analysis validates the biochemically defined NudE-dynein interaction domains *in vivo*, and a two-hit genetic approach reveals a reciprocal dependence between retrograde transport initiation and microtubule maintenance that governs the threshold for synaptic degeneration. Together, these findings offer mechanistic insights into how defects in dynein activation can produce the progressive, length-dependent neurodegeneration characteristic of dying-back neuropathies.

## Results

### Loss of *nudE* causes axon length-dependent synaptic degeneration

Microtubule integrity is essential for axonal transport and long-term neuronal maintenance. Mutations in *tubulin* genes and in cytoplasmic dynein motor complex components cause progressive neurodegeneration in vertebrate model systems and human patients (Lipka et al., 2013; Matamoros and Baas, 2016). To systematically identify microtubule-associated factors whose loss produces axon length-dependent degeneration in motoneurons, we screened microtubule regulatory components by motoneuron-specific RNAi at the larval NMJ. This screen identified *nudE*, the sole *Drosophila* ortholog of mammalian Nde1 and Ndel1 (Arthur et al., 2015; Bradshaw et al., 2013; Garrott et al., 2022; Wainman et al., 2009). In control animals, the NMJ is organized into a series of synaptic boutons, each containing multiple active zones where the presynaptic scaffold protein Bruchpilot (Brp) is in precise apposition to postsynaptic glutamate receptors (GluRIIC) (Figure 1A, F). In *nudE* mutant animals (*nudE^39A^*), we observed synaptic degeneration at distal boutons, defined by the loss of the presynaptic active zone marker Brp despite the continued presence of postsynaptic GluRIIC, accompanied by fragmentation of the presynaptic motoneuron membrane (Figure 1B, G; the membrane staining becomes discontinuous and accumulations of the membrane marker become apparent). Identical phenotypes were observed in *nudE^39A^* homozygous and *nudE^39A^*^/*Df*^ (*nudE^39A^*mutant allele over a complete deficiency (Df) of the *nudE* genomic locus) transheterozygous animals, confirming that the *nudE^39A^* allele is a genetic null (Wainman et al., 2009; Figure S1A). This phenotype represents presynaptic withdrawal with preserved postsynaptic structures, a hallmark of synaptic retraction and degeneration at the *Drosophila* NMJ (Eaton et al., 2002; Mushtaq et al., 2022; Pielage et al., 2005, 2008; Stephan et al., 2015; Valakh et al., 2012).

**Figure 1:**
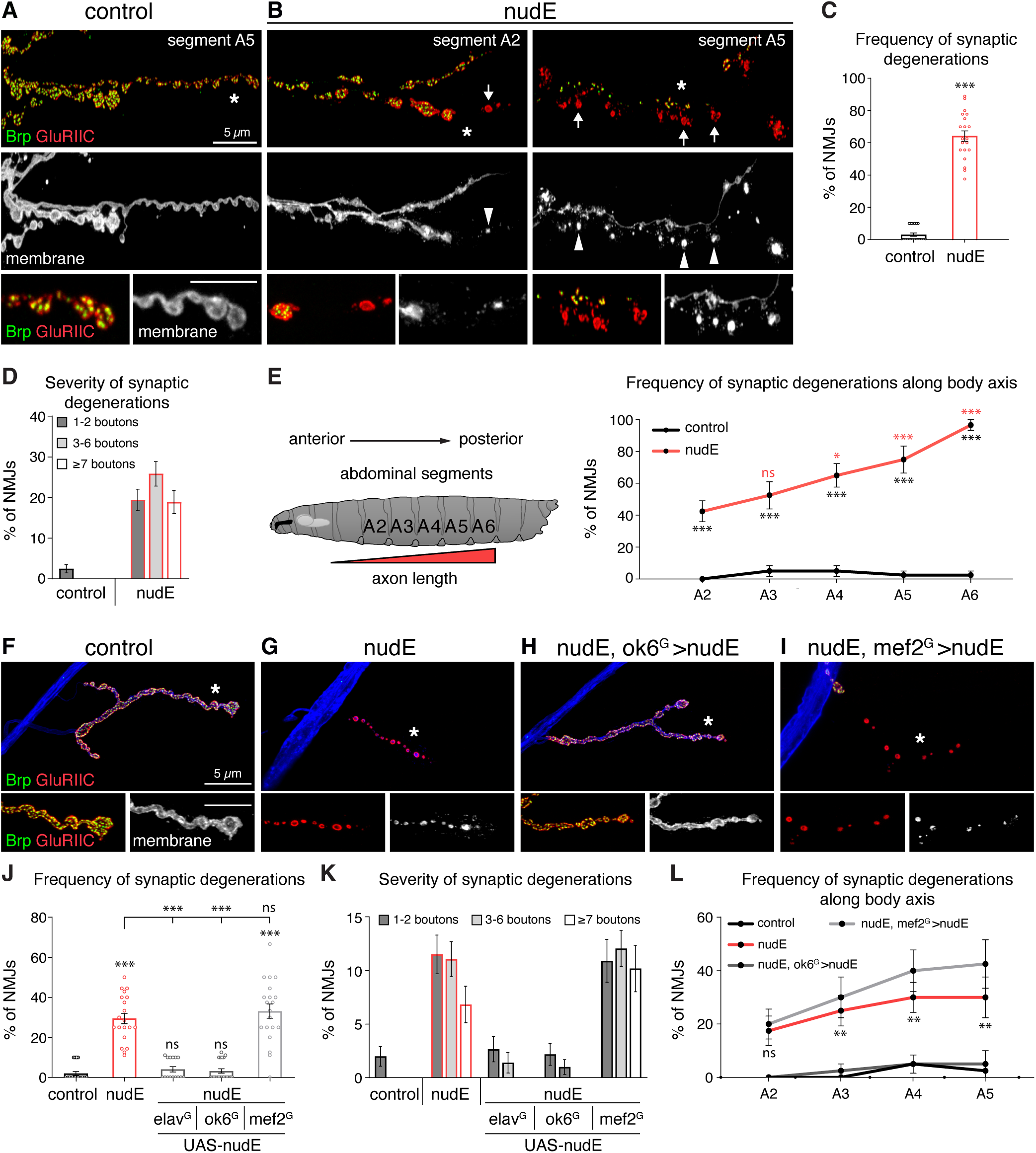
Loss of *nudE* causes axon length-dependent synaptic degeneration. (A, B) Confocal images of muscle 6/7 NMJs stained for the presynaptic active zone marker Brp (green), postsynaptic glutamate receptor GluRIIC (red), and neuronal membrane (HRP, gray). (A) In controls, Brp and GluRIIC are in precise apposition at all boutons. (B) In *nudE* mutants (*nudE^39A/39A^*), Brp is lost at individual boutons (arrows) while GluRIIC persists, and the neuronal membrane fragments (arrowheads). Degeneration is more severe at posterior segment A5 than at anterior segment A2. Asterisks mark regions shown at higher resolution in bottom panels. Scale bar, 5 µm. (C) Frequency of NMJs displaying synaptic degeneration at muscle 6/7. (D) Severity of degeneration, categorized by the number of affected boutons per NMJ. (E) Schematic of the larval body plan showing the anterior-to-posterior gradient of motoneuron axon length. Frequency of synaptic degeneration increases at posterior segments (A2–A6). (F–I) Confocal images of muscle 4 NMJs. (F) Control. (G) *nudE* mutant showing complete loss of presynaptic structures. (H) Motoneuron rescue (*nudE*, *ok6*-Gal4>UAS-*nudE*). (I) Muscle expression (*nudE*, *mef2*-Gal4>UAS-*nudE*) does not prevent degeneration. Asterisks mark regions shown at higher resolution in bottom panels. Scale bar, 5 µm. (J) Frequency of synaptic degeneration with tissue-specific rescue (elav^G^: pan-neuronal; ok6^G^: motoneuron; mef2^G^: muscle). (K) Severity of degeneration with tissue-specific rescue. (L) Frequency of degeneration along the body axis with motoneuron and muscle rescue. (C-E, J-L) Mean ± s.e.m. Welch’s t-test (C); two-way ANOVA with Sidak’s multiple comparisons test (E, L red stars indicate comparisons of A2 values with A3-A6 values for *nudE* mutants in E); Kruskal-Wallis with Dunn’s multiple comparisons test (J). ns, p ≥ 0.05; *p < 0.05; **p < 0.01; ***p < 0.001. Asterisks on top of columns denote comparisons with control, asterisks on horizontal lines comparisons between indicated genotypes. n = 20 animals per genotype (C–E, L); n = 20, 20, 15, 20, 20 animals (J, K).

Synaptic degeneration was highly penetrant, affecting more than 60% of NMJs at muscle 6/7 in *nudE* mutant animals (Figure 1C). Both the frequency and the severity of the phenotype increased along the anterior-posterior body axis. The proportion of NMJs displaying degeneration events rose from approximately 40% at segment A2 to nearly 100% at segment A6 (Figure 1D, E). At affected NMJs, the number of postsynaptic receptor fields lacking presynaptic Brp apposition, a measure of how many functional synapses are lost per NMJ, also increased in posterior segments (Figure 1D, S1E). In extreme cases, the entire NMJ degenerated with a complete loss of all presynaptic structures, a phenotype never observed in control animals (Figure 1G). This gradual increase in both frequency and severity with increasing motoneuron axon length was consistent across all muscle groups analyzed (muscles 1/9, 2/10, 4 and 6/7; Figure 1D, E, K, L, S1A–C, E) and resembles the distal-to-proximal progression characteristic of dying-back neuropathies in vertebrate model systems and human patients (Dadon-Nachum et al., 2011; Moloney et al., 2014; Rossor et al., 2012; Schaumburg et al., 1974).

All synaptic degeneration phenotypes were fully rescued by a bacterial artificial chromosome (BAC) encompassing the wild-type *nudE* genomic locus (Figure S1A–C). The BAC also rescued the pupal lethality of *nudE^39A^*mutant animals (Figure S1D), confirming that the *nudE^39A^* mutation is a clean loss-of-function allele and that the observed lethality is solely associated with the loss of *nudE*. Similarly, ubiquitous expression of a UAS-*nudE* transgene (*da*-Gal4) fully rescued lethality, verifying that this construct is sufficient to restore all essential NudE functions (Figure S1D). Pan-neuronal (*elav*-Gal4) and motoneuron-specific (*ok6*-Gal4) expression of the same transgene rescued synaptic degeneration and larval locomotion, whereas expression in postsynaptic muscles (*mef2*-Gal4) had no effect, demonstrating that NudE functions cell-autonomously in motoneurons to maintain synaptic stability (Figure 1F–L, S1A, B). However, motoneuron-specific expression did not rescue organismal viability (Figure S1D), indicating that NudE has additional essential functions outside the nervous system, in line with its known roles in cell division (Wainman et al., 2009).

### NudE is required for retrograde axonal transport initiation

We next addressed the cellular mechanism underlying synaptic degeneration in *nudE* mutants. NudE is an auxiliary subunit of the cytoplasmic dynein motor complex. *In vitro* reconstitution studies have established that the mammalian NudE ortholog Nde1 functions as a scaffold that tethers the regulatory factor Lis1 to autoinhibited dynein, enabling Lis1 to open the phi conformation and promote assembly of the processive dynein-dynactin-adaptor complex (Yang et al., 2026; Zhao et al., 2023; Zyłkiewicz et al., 2011). Nde1 is released upon dynactin binding and is not part of the active transport complex (Singh et al., 2024; Zhao et al., 2023). However, whether NudE/Nde1 is required specifically for transport initiation in neurons has not been directly tested.

To determine whether NudE loss impairs dynein motor complex function at the synapse, we analyzed the distribution of the retrograde motor protein Dynein heavy chain (Dhc) and the anterograde motor protein Kinesin heavy chain (Khc) at the NMJ. In control animals, neither Dhc nor Khc were detectable at synaptic boutons by confocal microscopy, reflecting an even distribution of individual motor complexes at low abundance and their rapid removal from synapses by retrograde transport (Figure 2A, S2A). In *nudE* mutants, both Dhc and Khc accumulated prominently at the distal-most synaptic bouton of more than 70% of all NMJs (Figure 2A, B; S2A, B). These accumulations were not restricted to motor proteins: membrane-associated glycoproteins (anti-HRP) (Figure 2A, S2B, 5A) and the synaptic vesicle transporter DvGlut (Figure 5A) showed similar accumulations at distal boutons indicating a general failure to clear retrograde cargo from the synaptic terminal. The co-accumulation of the anterograde motor Khc is expected, as Khc is itself a retrograde cargo that must be transported back toward the soma after delivering its anterograde payload (Lloyd et al., 2012). Analysis along the anterior-posterior body axis revealed that the proportion of NMJs with Khc accumulations followed the same distal-to-proximal gradient as synaptic degeneration, but was consistently higher at every segment examined (Figure S2B). All accumulation phenotypes were completely rescued by motoneuron expression of wild-type *nudE* (Figure 2A, B; S2B).

**Figure 2:**
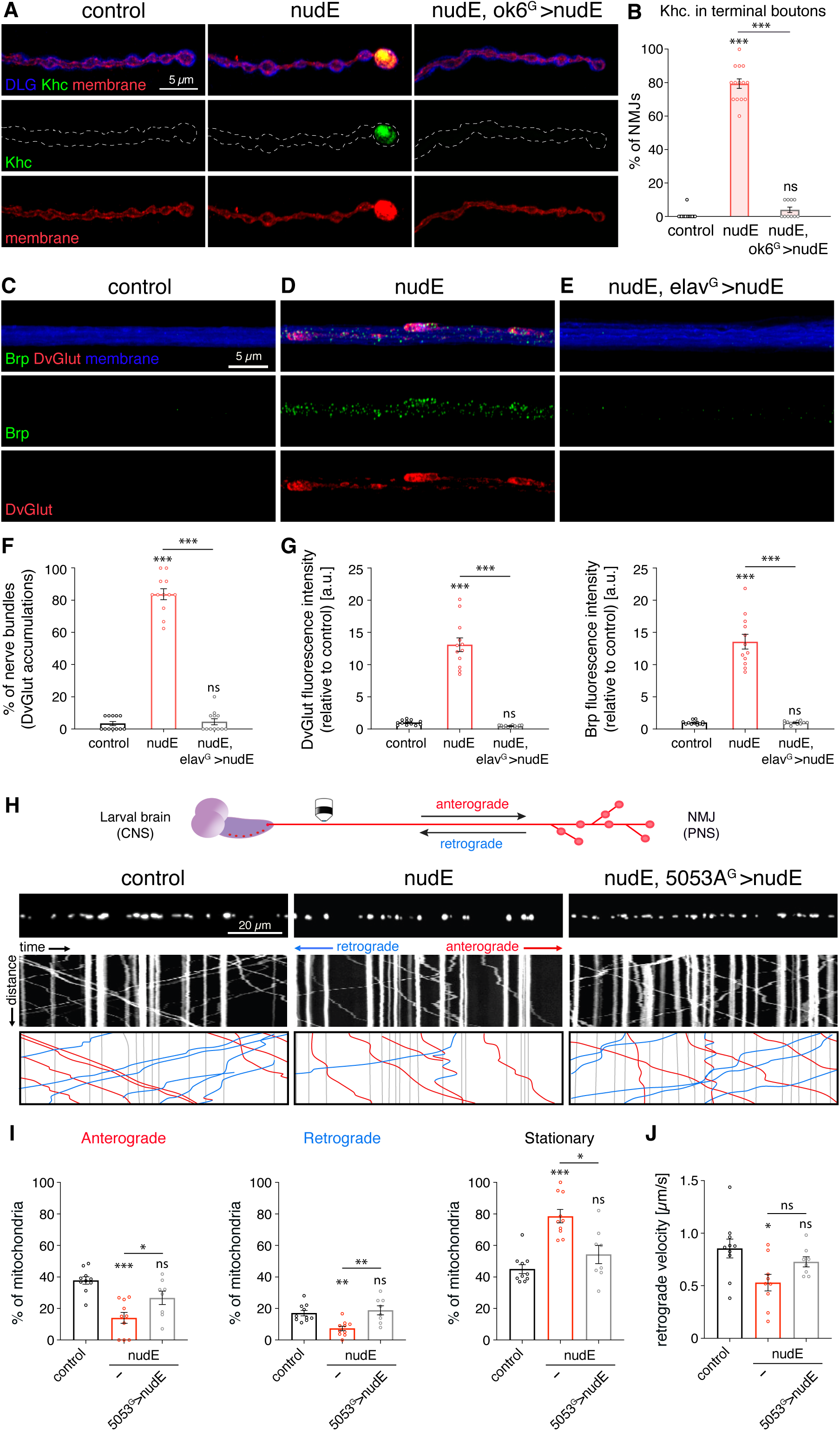
NudE is required for retrograde transport initiation and axonal transport. (A) Confocal images of muscle 4 NMJs stained for the postsynaptic marker Discs-large (DLG, a MAGUK family member, blue) to identify type Ib boutons, Kinesin heavy chain (Khc, green), and neuronal membrane (HRP, red). In controls, Khc is not detectable at synaptic boutons. In *nudE* mutants (*nudE^39A/39A^*), Khc and membrane-associated proteins accumulate at distal boutons. This phenotype is rescued by motoneuron expression of *nudE* (*nudE*, *ok6*-Gal4>UAS-*nudE*). Scale bar, 5 µm. (B) Frequency of NMJs with Khc accumulations at terminal boutons. (C–E) Confocal images of nerve bundles stained for the active zone protein Brp (green), synaptic vesicle transporter DvGlut (red), and neuronal membrane (HRP, blue). (C) In controls, only small puncta of Brp and DvGlut are detectable along nerve bundles. (D) In *nudE* mutants (*nudE^39A/39A^*), large accumulations of both markers are present. (E) Neuronal expression of *nudE* (*nudE*, *elav*-Gal4>UAS-*nudE*) rescues axonal transport defects. Scale bar, 5 µm. (F) Percentage of nerve bundles containing DvGlut aggregates. (G) Fluorescence intensity of DvGlut (left) and Brp (right) in nerve bundles, normalized to controls. (H) Top: Motoneuron schematic showing the direction of anterograde and retrograde transport between CNS and NMJ. Bottom: Representative images and kymographs of GFP-tagged mitochondria in individual motoneuron axons of control, *nudE* mutant, and *nudE* rescue (*nudE*, *5053A*-Gal4>UAS-*nudE*) animals. Anterograde transport is traced in red, retrograde in blue. Scale bar, 20 µm. (I) Quantification of the percentage of mitochondria per axon moving in the anterograde direction (left), retrograde direction (middle), or remaining stationary (right). (J) Retrograde velocity of motile mitochondria. (B, F, G, I, J) Mean ± s.e.m. Kruskal-Wallis with Dunn’s multiple comparisons test (B, F, I (stationary)); one-way ANOVA with Sidak’s multiple comparisons test (G, I anterograde and retrograde, J). ns, p ≥ 0.05; *p < 0.05; **p < 0.01; ***p < 0.001. n = 14, 14, 10 animals (B); n = 12 animals per genotype (F, G); n = 10, 10, 8 (anterograde); 9, 9, 14 (retrograde and stationary) axons from a minimum of 6 animals (I,); n = 10, 7, 8 axons from a minimum of 6 animals (J).

Consistent with a disruption of axonal transport, we observed large accumulations of the synaptic vesicle marker DvGlut and the active zone protein Brp in *nudE* mutant axon bundles, whereas only small puncta of these markers were detectable in controls (Figure 2C–E). Approximately 80% of *nudE* mutant nerve bundles contained DvGlut aggregates, and fluorescence intensities of both DvGlut and Brp were increased more than 10-fold compared to controls (Figure 2F, G). These accumulations were completely rescued by neuronal expression of wild-type *nudE* (Figure 2E–G).

The distal motor accumulations and axonal transport defects are consistent with two fundamentally different mechanisms: a general impairment of motor function during transport, or a specific failure to initiate transport at the synaptic terminal. To distinguish between these possibilities, we analyzed mitochondrial transport in motoneuron axons using live imaging (Figure 2H). In control axons, approximately 38% of mitochondria moved anterogradely, 17% retrogradely, and 45% were stationary, consistent with published values for *Drosophila* larval motor axons (Pilling et al., 2006).

In *nudE* mutants, the proportion of motile mitochondria was reduced in both directions, with a corresponding increase in stationary mitochondria to approximately 79% (Figure 2I). Total mitochondrial density was unchanged (Figure S2C), indicating that mitochondria are not lost but fail to initiate transport. Retrograde velocity of those mitochondria that did move was moderately reduced, while anterograde velocity was not significantly affected (Figure 2J, S2D), indicating a predominant impairment of dynein-dependent transport, with secondary effects on anterograde motility likely reflecting the dependence of kinesin on dynein-mediated recycling. Motoneuron-specific re-expression of NudE restored both transport initiation and retrograde velocity (Figure 2I, J). Thus, the predominant consequence of NudE loss is a reduction in the proportion of cargo entering transport, consistent with impaired dynein complex assembly at the synaptic terminal. The additional retrograde velocity reduction is in line with the known role of the NudE-Lis1 module in promoting the formation of active dynein complexes (Htet et al., 2020; Yang et al., 2026), which we next tested directly through structure-function analysis.

The distal motor accumulations in *nudE* mutants resemble the phenotype of a *Drosophila* model of the human motor neuron disease HMN7B, in which a G59S point mutation in the Cap-Gly domain of the dynactin subunit p150Glued (*Gl^G38S^*) causes Dhc and Khc accumulations at terminal boutons without affecting endosomal transport velocity along the axon (Lloyd et al., 2012). However, *Gl^G38S^* larvae did not display synaptic degeneration (Lloyd et al., 2012), whereas *nudE* loss progresses to structural degeneration with an additional reduction in retrograde velocity. This difference reflects the distinct molecular roles of the two proteins: the Cap-Gly domain mediates recruitment of dynactin to microtubule plus-ends, a step dispensable once transport has initiated (Moughamian and Holzbaur, 2012), whereas NudE acts on the motor itself by promoting active dynein complex formation (Yang et al., 2026; Zhao et al., 2023). Both proteins operate at the transport initiation step, but NudE loss has more severe consequences because it affects the assembly of active motor complexes upstream of motor-microtubule engagement.

Unlike dynactin, NudE does not directly bind microtubules but associates with the dynein motor complex through three defined interaction domains: a dynein intermediate chain (Dic) binding domain within the N-terminal coiled-coil, a Lis1-binding domain, and a dynein heavy chain (Dhc) binding domain in the C-terminal globular region (Figure 3A). Two independent studies identified a set of conserved acidic residues in the N-terminal coiled-coil of Ndel1/Nudel that mediate binding to the dynein intermediate chain through electrostatic interactions, and demonstrated that alanine substitution of these residues progressively weakens Dic binding (Wang and Zheng, 2011; Zyłkiewicz et al., 2011). This interaction surface was subsequently confirmed as a conserved and essential hub for dynein activation (Okada et al., 2023). These residues are conserved between *Drosophila* NudE and both human paralogs Nde1 and Ndel1 (Figure 3A).

**Figure 3:**
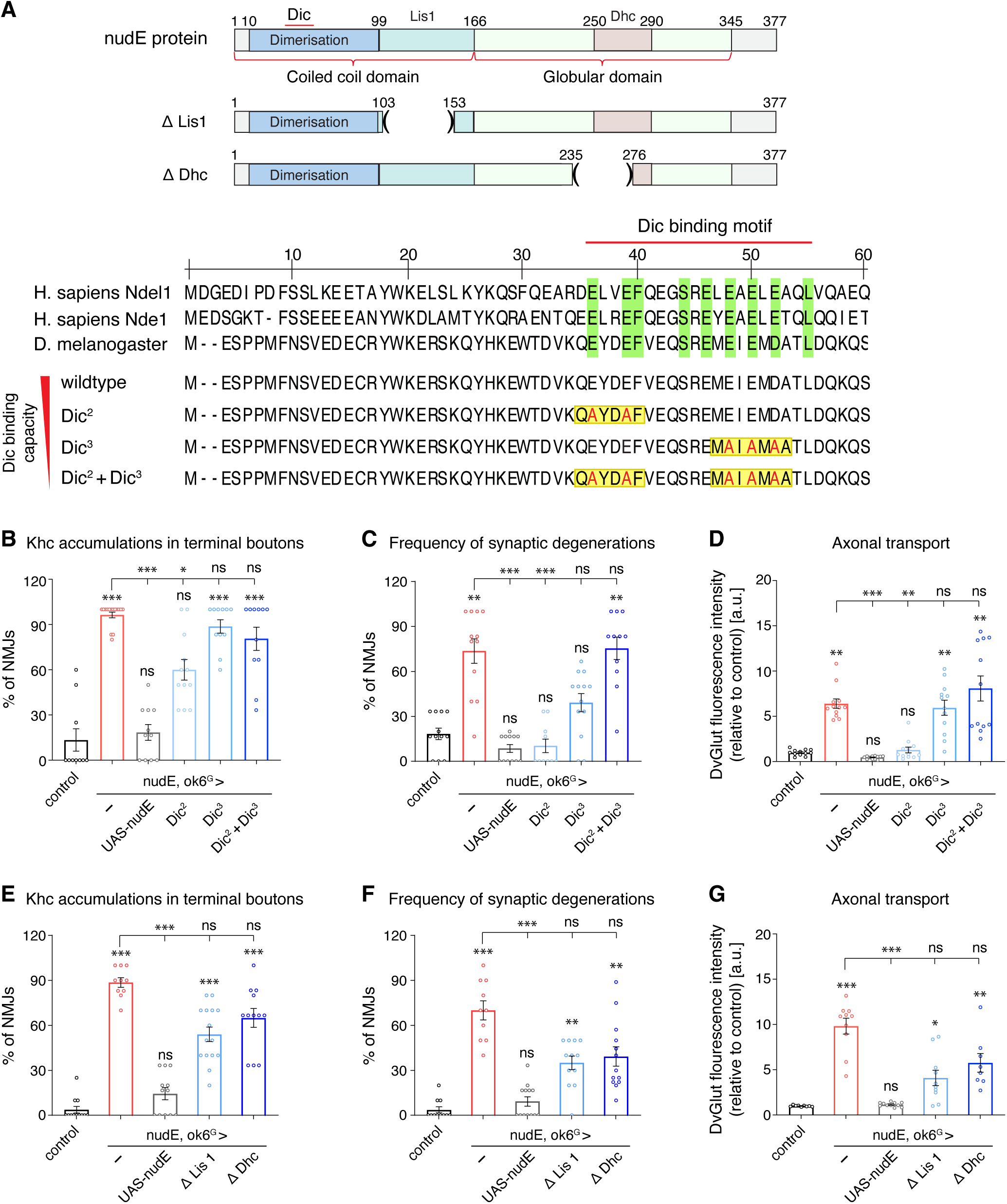
Structure-function analysis validates the NudE-dynein interaction domains *in vivo*. (A) Top: Domain structure of the NudE protein showing the Dic-, Lis1-, and Dhc-binding domains, and schematic of *nudE* constructs lacking the Lis1-binding (*ΔLis1*, Δ103–153) or Dhc-binding (*ΔDhc*, Δ235–276) domain. Middle: Alignment of the Dic-binding motif of human Ndel1, Nde1, and *Drosophila* NudE with conserved residues highlighted in green. Bottom: The *nudE^Dic2^*construct carries two glutamate-to-alanine substitutions (E34A/E37A) targeting the milder of two biochemically defined binding sites, *nudE^Dic3^*carries three substitutions (E46A/E48A/D50A) targeting the more critical site, and *nudE^Dic2+Dic3^* combines all five mutations. These mutations progressively reduce binding to the dynein intermediate chain. (B–D) Quantification of Khc accumulations at terminal boutons (B, muscle 6/7), frequency of synaptic degeneration (C, muscle 6/7), and axonal transport measured by DvGlut fluorescence intensity in nerve bundles (D) for wild-type and Dic-binding mutant constructs expressed in motoneurons of *nudE* mutants. (E-G) Quantification of Khc accumulations, synaptic degeneration, and axonal transport for *nudE* constructs lacking the Lis1-binding (*ΔLis1*, Δ103–153) or Dhc-binding (*ΔDhc*, Δ235–276) domain expressed in motoneurons of *nudE* mutants. (B-G) Mean ± s.e.m. Kruskal-Wallis with Dunn’s multiple comparisons test (B-G). ns, p ≥ 0.05; *p < 0.05; **p < 0.01; ***p < 0.001. n = 10, 15, 11, 12, 11, 11 animals (B); n = 12, 12, 12, 10, 13, 11 animals (C); n = 12 animals per genotype (D); n = 12, 10, 12, 15, 12 animals (E); n = 11, 10, 13, 12, 13 animals (F); n = 9, 10, 11, 10, 8 animals (G).

To test whether the NudE-Dic interaction is required for retrograde transport initiation *in vivo*, we generated three transgenic constructs carrying progressively more severe mutations in the Dic-binding domain: *nudE^Dic2^* (E34A/E37A), corresponding to the milder of the two biochemically characterized binding sites; *nudE^Dic3^* (E46A/E48A/D50A), targeting the more critical binding site; and the combined *nudE^Dic2+Dic3^*(E34A/E37A/E46A/E48A/D50A), predicted to abolish Dic binding entirely (Figure 3A). All constructs were integrated at the same genomic locus (*attP40*) to ensure comparable expression levels. We expressed wild-type or mutant NudE in motoneurons of *nudE* mutant animals and assessed three phenotypic parameters: Khc accumulations in terminal boutons, frequency of synaptic degeneration, and axonal transport as measured by DvGlut fluorescence intensity in nerve bundles.

Wild-type NudE efficiently rescued all three parameters (Figure 3B–D). The mutant constructs showed a graded rescue that correlated with predicted residual Dic-binding capacity. The mildest mutation (*nudE^Dic2^*) completely rescued synaptic degeneration and axonal transport deficits but only partially reduced Khc accumulations in terminal boutons. The more severe *nudE^Dic3^*partially rescued synaptic degeneration but failed to significantly improve axonal transport or Khc accumulations. The combined *nudE^Dic2+Dic3^*mutant failed to rescue any parameter (Figure 3B–D). This graded *in vivo* rescue directly parallels the biochemical findings: Zyłkiewicz et al. (2011) showed that the equivalent Ndel1 E36A/E39A mutation (corresponding to *nudE^Dic2^*E34A/E37A) retained 70% rescue activity in a microtubule aster formation assay, whereas Ndel1 E48A/E52A (corresponding to *nudE^Dic3^* E46A/E48A/D50A) largely abolished function. Zhao et al. (2023) subsequently demonstrated that the corresponding Nde1-E47A point mutation (equivalent to Ndel1 E48A and NudE E48A within the Dic3 site) completely prevented Nde1 from promoting Lis1-mediated dynein activation in reconstituted motility assays. Our *in vivo* data thus validate the biochemically defined Dic-binding interface as essential for NudE function in neurons and establish that the graded disruption of this interface produces correspondingly graded transport initiation and synaptic maintenance defects.

To determine whether the Lis1 and Dhc binding domains of NudE are similarly essential, we generated rescue constructs lacking either the previously characterized Lis1-binding region (*nudE^ΔLis1^*, Δ103–153; (Derewenda et al., 2007)) or the Dhc-binding region (*nudE^ΔDhc^*, Δ235–276; (Sasaki et al., 2000)) (Figure 3A, E-G). Neither deletion construct significantly rescued any of the three parameters, although both showed a consistent trend toward partial improvement (Figure 3E-G). Thus, NudE function requires all three interaction domains, Dic, Lis1, and Dhc, as expected for its role as a scaffold that must simultaneously engage the dynein motor and its regulatory partner Lis1 to promote transport complex assembly (Derewenda et al., 2007; Zyłkiewicz et al., 2011).

The structure-function analysis predicts that perturbation of any component of the dynein motor complex should phenocopy *nudE* loss, because the assembly of an active transport complex requires the stoichiometric engagement of multiple dynein and dynactin subunits (Garrott et al., 2022). To test this directly, we knocked down six additional dynein-dynactin complex components in motoneurons via RNAi or dominant-negative expression: *Dic*, *Dlic*, *Dhc*, *Dynamitin* (*Dmn*), *Dynactin*/*p150^Glued^*(*Gl^DN^*), and *Lis1* (Figure S3). In every case, we observed severe Khc accumulations at distal boutons, significant increases in synaptic degeneration, and large accumulations of DvGlut and Brp in axon bundles (Figure S3I, J, K–R). Synaptic retractions were previously reported for loss of the *dynactin* subunit *p150^Glued^* (Eaton et al., 2002); our data demonstrate that perturbation of any component of the dynein-dynactin complex produces a similar phenotype. Importantly, in all genotypes the frequency of Khc accumulations at terminal boutons exceeded the frequency of synaptic degeneration (Figure S3I, J). This consistent hierarchy, with transport initiation defects more frequent than synaptic degeneration across all perturbations of the dynein-dynactin complex, indicates that the failure to initiate retrograde transport precedes and likely causes the subsequent loss of synaptic stability.

### Loss of *nudE* induces progressive neurodegeneration

To determine whether the synaptic degeneration observed in *nudE* mutants is progressive or reflects a developmental failure, we performed non-invasive live imaging of NMJs in intact, behaving larvae over a 48-hour period spanning the second to third larval instar stage. For this, we co-expressed fluorescently labeled active zone (*Brp-mStraw*) and membrane (*mCD8-GFP*) markers in motoneurons of control and *nudE* mutant animals and imaged the same NMJs every 12 hours (Figure 4A, B). In both control and *nudE* mutant larvae, NMJs grew normally during this period, with new synaptic boutons being added at comparable rates. This confirms that NudE is not required for synapse formation or growth, and that the degeneration phenotype represents a failure of synapse maintenance rather than a developmental defect. In *nudE* mutants, the first signs of degeneration appeared at 12 hours, when membrane accumulations became detectable at terminal boutons (Figure 4A, B). Over the following 24–36 hours, these accumulations increased in size and became evident at all terminal boutons. By 48 hours, the Brp-mStraw signal at affected boutons began to dissolve, indicating disassembly of presynaptic active zones (Figure 1B, G; Figure 4A, B). Thus, degeneration in *nudE* mutants is progressive, initiates at the distal-most boutons, and follows a stereotyped sequence: membrane and cargo accumulations at terminal boutons precede the loss of presynaptic active zone organization.

**Figure 4:**
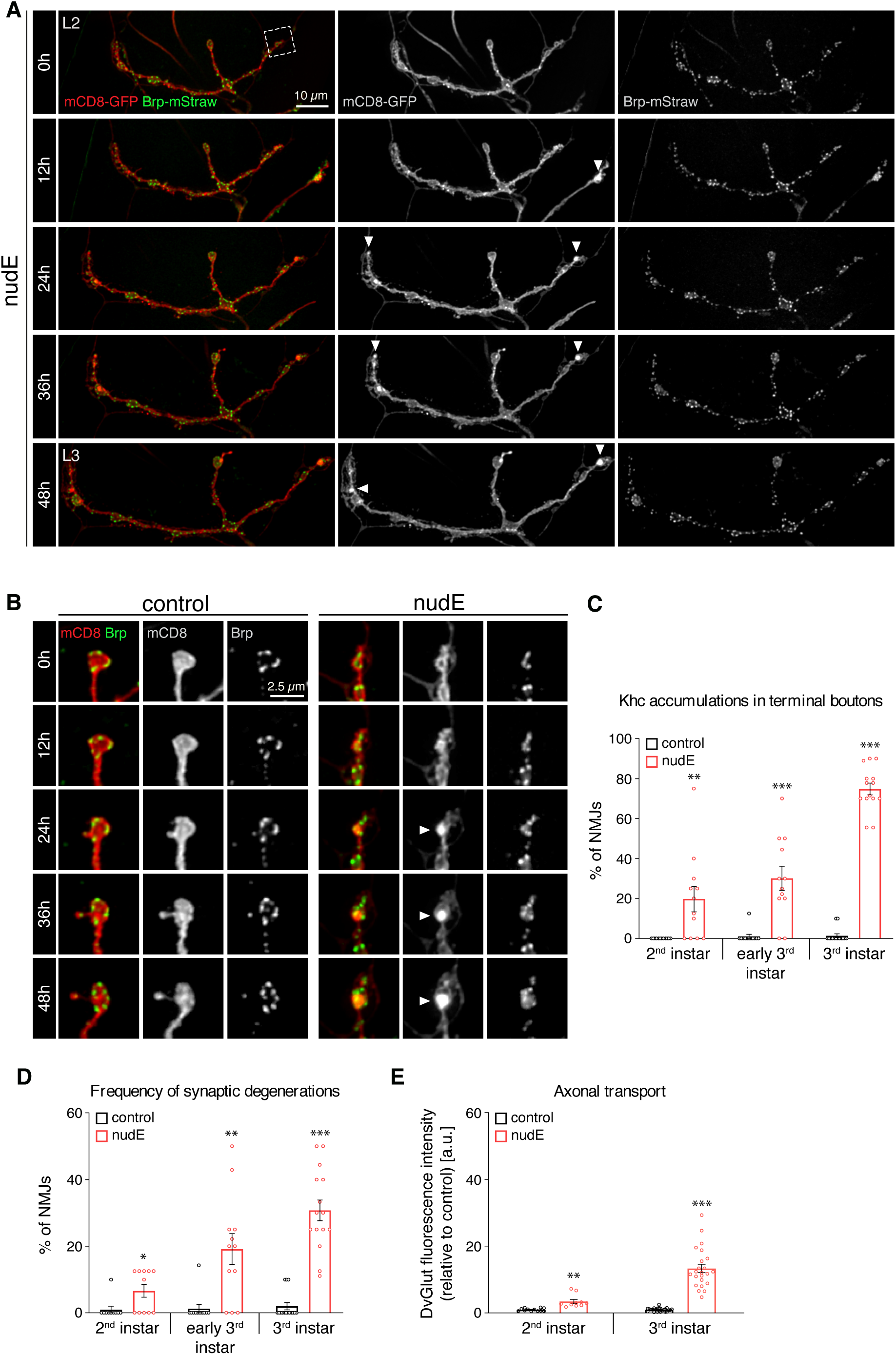
Transport initiation failure precedes synaptic degeneration and axonal transport defects. (A) Non-invasive live imaging of NMJ development and degeneration over 48 h spanning the second to third larval instar. Motoneuron membranes are labeled with mCD8-GFP (red) and presynaptic active zones with Brp-mStraw (green), shown individually in gray. At 0 h, *nudE* mutant (*nudE^39A/39A^*) NMJs are normally developed. From 12 h onward, membrane accumulations appear at terminal boutons (arrowheads) and progressively increase. NMJs continue to grow and add new boutons throughout this period in both control and *nudE* mutant animals. Scale bar, 10 µm. (B) Higher magnification of terminal boutons from control and *nudE* mutant NMJs at each time point. Arrowheads indicate progressive membrane accumulations at terminal boutons; at these sites, Brp puncta dissolve over time. Scale bar, 2.5 µm. (C–E) Quantification of Khc accumulations at terminal boutons (C), frequency of synaptic degenerations (D), and axonal transport measured by DvGlut fluorescence intensity in nerve bundles (E) at the indicated developmental stages. Khc accumulations are the earliest detectable defect, present from the 2^nd^ instar stage and increasing over time. Synaptic degeneration and axonal transport defects increase progressively, with the most pronounced differences at the 3^rd^ instar stage. (C-E) Mean ± s.e.m.. Mann Whitney test (C-E); Welch’s test (E, 2^nd^ instar). ns, p ≥ 0.05; *p < 0.05; **p < 0.01; ***p < 0.001. n = 12, 12, 14, 12, 12, 14 animals (C); n = 10, 11, 15, 11, 12, 15 animals (D); n = 12, 24, 10, 23 animals (E).

To put the subcellular progression of degeneration observed by live imaging into a more quantitative framework and compare it to the temporal sequence of axonal transport defects, we analyzed three parameters using fixed preparations: Khc accumulations in terminal boutons, frequency of synaptic degeneration, and axonal transport defects, at three developmental stages: second instar, early third instar, and third instar (Figure 4C–E). Khc accumulations at terminal boutons were the earliest detectable defect, already present at 19.8±6.4% of NMJs in second instar *nudE* mutant larvae and rising to 74.8±2.9% by third instar (Figure 4C). In contrast, only minor synaptic degeneration defects could be observed at second instar NMJs (6.6±1.9%), but these values progressively increased to 19.2±4.6% and 30.8±3.1% during the third instar stages (Figure 4D). Axonal transport defects, measured as DvGlut fluorescence intensity in nerve bundles, were hardly detectable at the second instar stage and only became highly significant at the third instar (Figure 4E). This temporal hierarchy, transport initiation failure first, followed by synaptic degeneration, with axonal transport disruption appearing last, is consistent with the observation that the frequency of Khc accumulations significantly exceeds the frequency of synaptic degeneration across all dynein-dynactin complex perturbations (Figure S3I, J). Together, these data establish that the failure to initiate retrograde transport is the primary defect in *nudE* mutant motoneurons, from which the loss of synaptic stability progressively develops.

### *nudE* is required for the maintenance of the synaptic microtubule cytoskeleton

To address how deficits in retrograde transport initiation lead to synaptic degeneration, we next analyzed the presynaptic microtubule cytoskeleton. In control animals, microtubules extend continuously through all synaptic boutons and become gradually thinner toward the distal end of the NMJ, as visualized by staining for Futsch, the *Drosophila* homolog of the microtubule-associated protein MAP1B (Figure 5A). In *nudE* mutant animals, we observed a variable disruption of this pattern. Phenotypes ranged from NMJs with minor perturbations of the terminal microtubule cytoskeleton accompanied by DvGlut accumulations in terminal boutons, to NMJs with multiple gaps in the Futsch staining but an intact presynaptic membrane, to NMJs with severe Futsch gaps and fragmentation of the presynaptic membrane (Figure 5A). Quantification revealed that 37.5±2.5% of NMJs displayed gaps in Futsch staining, whereas 85.0±4.2% showed DvGlut accumulations in terminal boutons (Figure 5B, C). The significantly higher frequency of DvGlut accumulations compared to Futsch gaps indicates that cargo accumulations at terminal boutons, a consequence of failed retrograde transport initiation, precede the disruption of the microtubule cytoskeleton. All phenotypes were rescued by motoneuron expression of wild-type *nudE* (Figure 5B, C).

**Figure 5:**
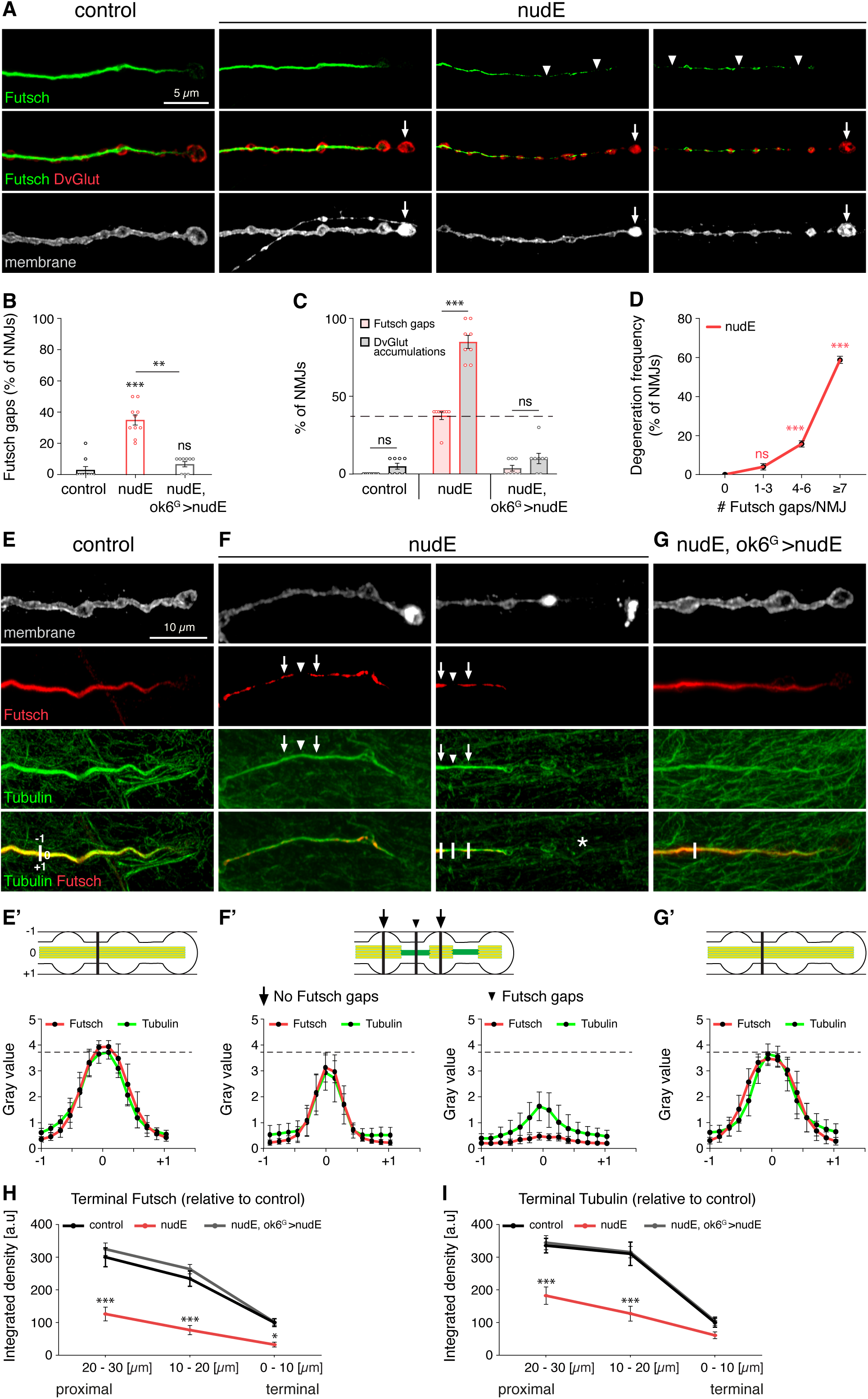
*nudE* is required for the maintenance of the synaptic microtubule cytoskeleton. (A) Confocal images of muscle 4 NMJs stained for the microtubule-associated protein Futsch/MAP1B (green), synaptic vesicle transporter DvGlut (red), and neuronal membrane (HRP, gray). In controls, Futsch forms a continuous structure that gradually thins toward terminal boutons (left). In *nudE* mutants (*nudE^39A/39A^*), phenotypes range from mild DvGlut and membrane accumulations at terminal boutons with intact Futsch (left, arrows), to multiple gaps in Futsch distribution (middle, arrowheads), to severe disruptions with fragmentation of the motoneuron membrane (right). Scale bar, 5 µm. (B) Percentage of muscle 4 NMJs displaying Futsch gaps. (C) Frequency of Futsch gaps compared to DvGlut accumulations. DvGlut accumulations are significantly more frequent than Futsch gaps, indicating that cargo accumulations at terminal boutons precede microtubule destabilization. (D) Frequency of synaptic degeneration as a function of the number of Futsch gaps per NMJ. The likelihood of degeneration increases progressively with the number of Futsch gaps. (E–G’) Analysis of Tubulin levels (green) in relation to Futsch (red) and neuronal membrane (HRP, gray). Line scans (E’, F’, G’) show fluorescence intensity profiles across the NMJ with Futsch gaps (arrowheads) and sites without gaps (arrows) indicated. (E, E’) In controls, Futsch and Tubulin are present throughout the terminal. (F, F’) In *nudE* mutants, regions retaining Futsch contain near-normal Tubulin levels (arrows). At Futsch-negative sites (arrowheads), Tubulin is reduced but still present. At sites of membrane fragmentation (asterisk), both markers are absent. (G, G’) Motoneuron rescue (*nudE*, *ok6*-Gal4>UAS-*nudE*). Scale bar, 10 µm. (H, I) Integrated density of Futsch (H) and Tubulin (I) within the last 30 µm of the NMJ, measured in 10 µm intervals from the tip of the terminal bouton. Both markers are severely reduced in *nudE* mutants, with the strongest reduction at the distal-most 0–10 µm. (B-D, É-I) Mean ± s.e.m. Kruskal-Wallis with Dunn’s multiple comparisons test (B); two-way ANOVA with Tukey’s multiple comparisons test (C, H, I); one-way ANOVA with Sidak’s multiple comparison tests (D, red stars indicate comparisons to the zero (0) Futsch gaps/NMJ condition). ns, p ≥ 0.05; *p < 0.05; **p < 0.01; ***p < 0.001. n = 10, 10, 9 animals (B); n = 8 animals per genotype (C); n = 7, 8, 7, 14 animals (D); n = 46, 25, 41 NMJs from 8 animals per genotype (E–G’); n = 20, 29, 20 NMJs from 8 animals per genotype (H); n = 20, 26, 20 NMJs from 8 animals (I).

To determine whether the extent of microtubule disruption correlates with the likelihood of synaptic degeneration, we quantified the frequency of degeneration as a function of the number of Futsch gaps per NMJ in *nudE* mutants. NMJs without Futsch gaps did not display synaptic degeneration, whereas the frequency of degeneration increased progressively with the number of gaps: from 4.0±1.7% at NMJs with 1–3 gaps to 15.7±1.6% with 4–6 gaps and 58.8±1.9% at NMJs with seven or more gaps (Figure 5D). This indicates that progressive disruption of the microtubule cytoskeleton increases the vulnerability of the synapse to degeneration.

Futsch/MAP1B is a microtubule-associated protein that preferentially binds to bundled microtubules. The absence of Futsch staining therefore does not necessarily indicate a complete loss of microtubules. To directly assess microtubule integrity, we co-stained *nudE* mutant NMJs for Futsch and Tubulin. In control animals, both markers colocalized throughout the nerve terminal (Figure 5E, E’). In *nudE* mutants, regions that retained Futsch staining contained near-normal levels of Tubulin (Figure 5F, F’). At sites where Futsch was absent, Tubulin-positive microtubules were still present but at significantly reduced levels, indicating that the microtubule cytoskeleton is diminished but not eliminated at these positions (Figure 5F, F’). In contrast, at sites of presynaptic membrane fragmentation, both Futsch and Tubulin were absent (Figure 5F, asterisk). To quantify this gradient in an unbiased manner, we measured the integrated density of both Futsch and Tubulin within the last 30 µm of the NMJ without prior classification of individual boutons. In *nudE* mutants, both Futsch and Tubulin levels were severely reduced in the terminal region, with the most pronounced loss at the distal-most 0–10 µm (Futsch: 32.7±7.3% of control; Tubulin: 61.3±10.4% of control) (Figure 5H, I). All microtubule phenotypes were rescued by expression of wild-type *nudE* in motoneurons (Figure 5E–I). These data indicate that loss of NudE leads to a progressive reduction of microtubule levels at distal boutons, with Futsch dissociating from microtubules as bundling decreases below a critical threshold. The complete loss of both Futsch and Tubulin occurs only at sites that have progressed to membrane fragmentation and synaptic degeneration.

### NudE is required to maintain synaptic transmission

To determine whether the observed structural defects directly impair neuronal function, we performed electrophysiological recordings at NMJs in two segments along the anterior-posterior body axis that are differentially affected by synaptic degeneration: segment A3 (proximal, mildly affected) and segment A6 (distal, severely affected). At segment A3, miniature excitatory junctional potential (mEJP) amplitude (0.89±0.06 mV vs 0.86±0.05 mV in controls), evoked excitatory junctional potential (EJP) amplitude (26.8±1.3 mV vs 28.9±2.6 mV), and quantal content (31.8±2.0 vs 33.3±1.7) were not significantly different between *nudE* mutants and controls (Figure 6A, A’; Figure S4A–C). In contrast, at segment A6, *nudE* mutants displayed unchanged mEJP amplitude (0.70±0.05 mV vs 0.83±0.04 mV), but a significant reduction in EJP amplitude (14.1±1.3 mV vs 25.4±2.0 mV, a 45% decrease), and quantal content (20.4±1.3 vs 30.9±2.4, a 34% decrease) (Figure 6B, B’; Figure S4D–F). All parameters were rescued by motoneuron expression of wild-type *nudE* (Figure 6A’, B’; Figure S4). This functional gradient directly mirrors the structural degeneration gradient and confirms that the loss of presynaptic active zones at distal NMJs results in a corresponding impairment of synaptic transmission.

**Figure 6:**
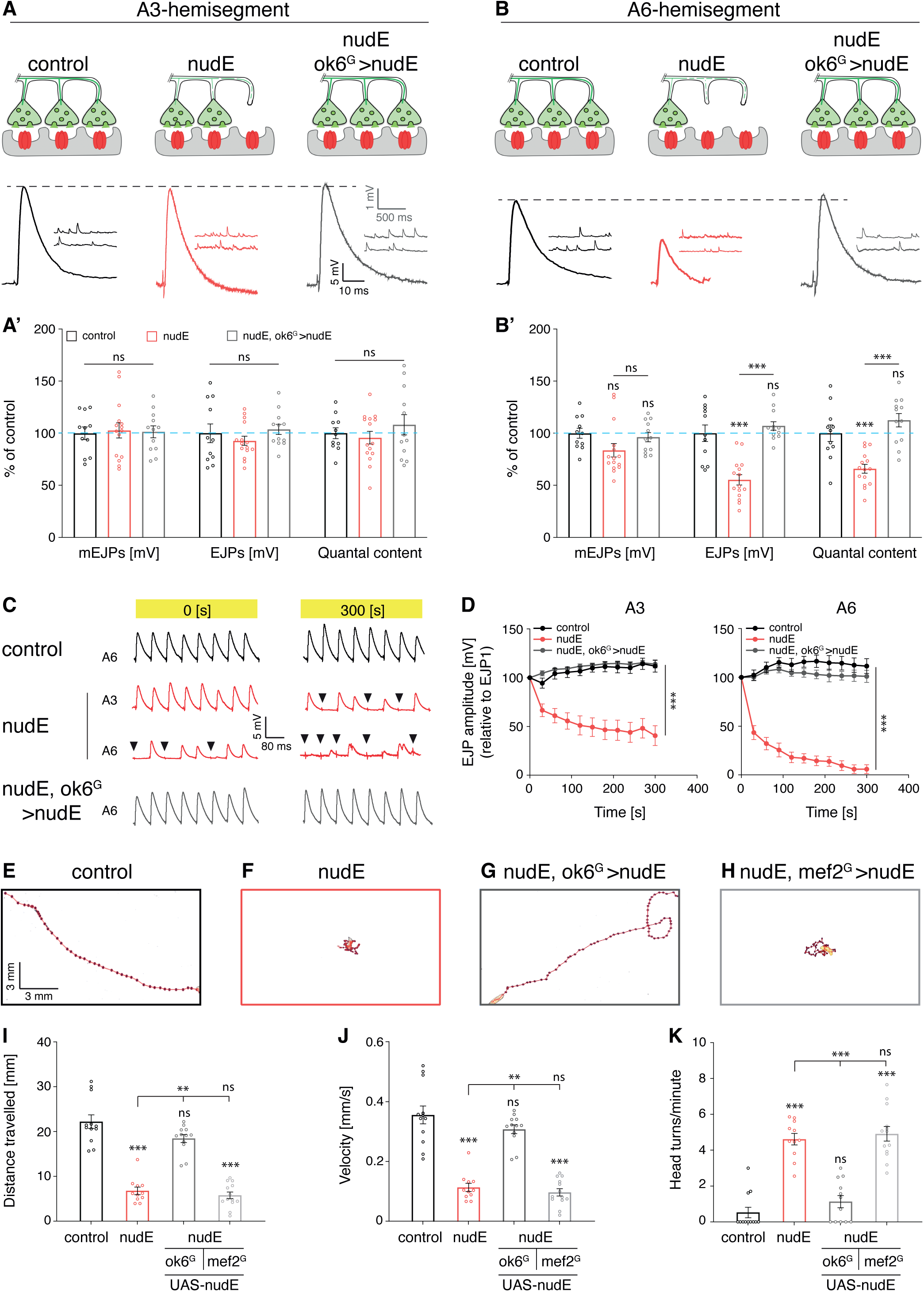
NudE is required to maintain synaptic transmission and locomotion. (A, B) Electrophysiological recordings at muscle 6/7 NMJs in proximal segment A3 (A) and distal segment A6 (B). Top: Schematics illustrate the extent of synaptic degeneration at each segment. Bottom: Representative traces of spontaneous miniature excitatory junctional potentials (mEJPs) and evoked excitatory junctional potentials (EJPs) are shown for control, *nudE* mutants (*nudE^39A/39A^*), and motoneuron rescue (*nudE*, *ok6*-Gal4>UAS-*nudE*). (A’, B’) Quantification of mEJP amplitude, EJP amplitude, and quantal content. At segment A3, no significant differences are observed between *nudE* mutants and controls. At segment A6, mEJP amplitude remains unchanged but EJP amplitude, and quantal content are significantly reduced in *nudE* mutants. All parameters are rescued by motoneuron expression of *nudE*. (C, D) Repetitive stimulation at 10 Hz for 5 min. (C) Representative traces showing responses to the initial and final stimuli. In controls, transmission is sustained at both segments. In *nudE* mutants, transmission failures (arrowheads) are observed from the onset at A6 and toward the end of the stimulation train at A3. (D) EJP amplitudes normalized to the first stimulus over the stimulation period. *nudE* mutants show rapid rundown at both A3 and A6, with more severe decline at A6. All deficits are rescued by motoneuron expression of *nudE*. (E–H) Representative locomotion trajectories of control (E), *nudE* mutant (F), motoneuron rescue (G), and muscle rescue (H) larvae. (I–K) Quantification of distance traveled per minute (I), velocity (J), and head turns per minute (K). *nudE* mutants display severely reduced locomotion that is rescued by motoneuron (*ok6*-Gal4) but not muscle (*mef2*-Gal4) expression of *nudE*. (Á, B’, D, I-K) Mean ± s.e.m. one-way ANOVA with Sidak’s multiple comparisons test (A’-B’); two-way ANOVA with Tukey’s multiple comparisons test (D); Kruskal-Wallis test with Dunn’s multiple comparisons test (I-K). ns, p ≥ 0.05; *p < 0.05; **p < 0.01; ***p < 0.001. n = 11, 15, 12 animals (A’, B’); n = 14, 17, 14 animals (D); n = 12, 11, 12, 13 animals (I–K).

To assess the functional capacity of the synapse under sustained activity, we applied repetitive stimulation at 10 Hz for 5 minutes and monitored EJP amplitudes over time (Figure 6C, D). In control animals, EJP amplitudes remained stable throughout the stimulation period at both A3 and A6. In *nudE* mutants, EJP amplitudes declined rapidly at both segments, dropping to approximately 40% of the initial amplitude at A3 and to approximately 18% at A6 by the end of the stimulation period (Figure 6D). In addition, *nudE* mutants displayed frequent failure events in which no vesicles were released in response to a stimulus. At A6, failures were already observed at the onset of the stimulation protocol and became more frequent over time. At A3, failures became evident toward the end of the stimulation train (Figure 6C). Such failures were never observed in control animals under these conditions. The pronounced rundown and failure events at A6 indicate a severely depleted synaptic vesicle pool at distal NMJs. Even at segment A3, where baseline transmission is normal, the inability to sustain transmission under repetitive stimulation indicates that the functional reserve of the synapse is already compromised at proximal segments with minimal structural degeneration. All transmission deficits were fully rescued by motoneuron expression of wild-type *nudE* (Figure 6D).

Consistent with the impairment of synaptic transmission, *nudE* mutant larvae displayed severe locomotion defects. Total distance traveled was reduced to 6.8±0.8 mm compared to 22.2±1.5 mm in controls, and velocity dropped from 0.36±0.03 to 0.11±0.01 mm/s (Figure 6E–F, I, J). In addition, *nudE* mutant larvae displayed a marked increase in head turns (4.6±0.3 per minute vs 0.8±0.4 in controls) and a characteristic tail-flip phenotype in which muscle contractions could not be properly executed in posterior segments (Figure 6F, K). This locomotion phenotype correlates with the posterior bias of both structural degeneration and synaptic transmission deficits and demonstrates that the cellular defects observed at individual NMJs have a direct functional consequence at the organismal level. All locomotion phenotypes were rescued by motoneuron (*ok6*-Gal4) but not muscle (*mef2*-Gal4) expression of wild-type *nudE* (Figure 6G, H, I–K).

### Progressive neurodegeneration in a two-hit model

Our data establish that defects in retrograde axonal transport initiation precede the disruption of the presynaptic microtubule cytoskeleton, which in turn increases the vulnerability of the synapse to degeneration. However, there appear to be thresholds in this process: we identified conditions in which transport initiation defects were present but did not lead to synaptic degeneration, including the *nudE^Dic2^* rescue (Figure 3B–D) and the *Gl^G38S^* disease model (Figure 7C, D). To test whether combining two sub-threshold perturbations can synergistically trigger degeneration, we generated double mutants between the Dynactin *Gl^G38S^* mutation, which disrupts retrograde transport initiation (Lloyd et al., 2012), and a *futsch*/*MAP1B* null mutation (*futsch^K68^*), which causes minor perturbations of the presynaptic microtubule cytoskeleton without affecting synaptic stability (Roos et al., 2000; Stephan et al., 2015). In *futsch^K68^*single mutants, we observed a moderate increase in Khc accumulations at terminal boutons (38.7±5.1%) that was not accompanied by HRP accumulations, indicating a weaker perturbation of retrograde transport initiation than in *Gl^G38S^*mutants (67.7±2.9% Khc with concurrent HRP accumulations) (Figure 7A, C). Neither *futsch^K68^* (4.0±1.6%) nor *Gl^G38S^* (1.8±1.2%) single mutants displayed synaptic degeneration (Figure 7D). In striking contrast, *futsch^K68^*; *Gl^G38S^* double mutants showed a significant increase in synaptic degeneration (23.6±3.1%), accompanied by a further increase in Khc accumulations (80.0±3.4%), disruption of microtubule integrity at terminal boutons, and fragmentation of the presynaptic membrane (Figure 7A–D). Analysis of Tubulin levels revealed that while single mutants showed only minor reductions of microtubule density restricted to the distal-most 10 µm of the NMJ, the double mutant displayed a significant reduction across the entire terminal region (Figure 7F, G). Consistent with the synaptic phenotypes, locomotion was only mildly affected in either single mutant but severely impaired in the double mutant (0.14±0.01 mm/s vs 0.42±0.03 mm/s in controls) (Figure 7H). Axonal transport was not significantly affected in any genotype (Figure 7E).

**Figure 7:**
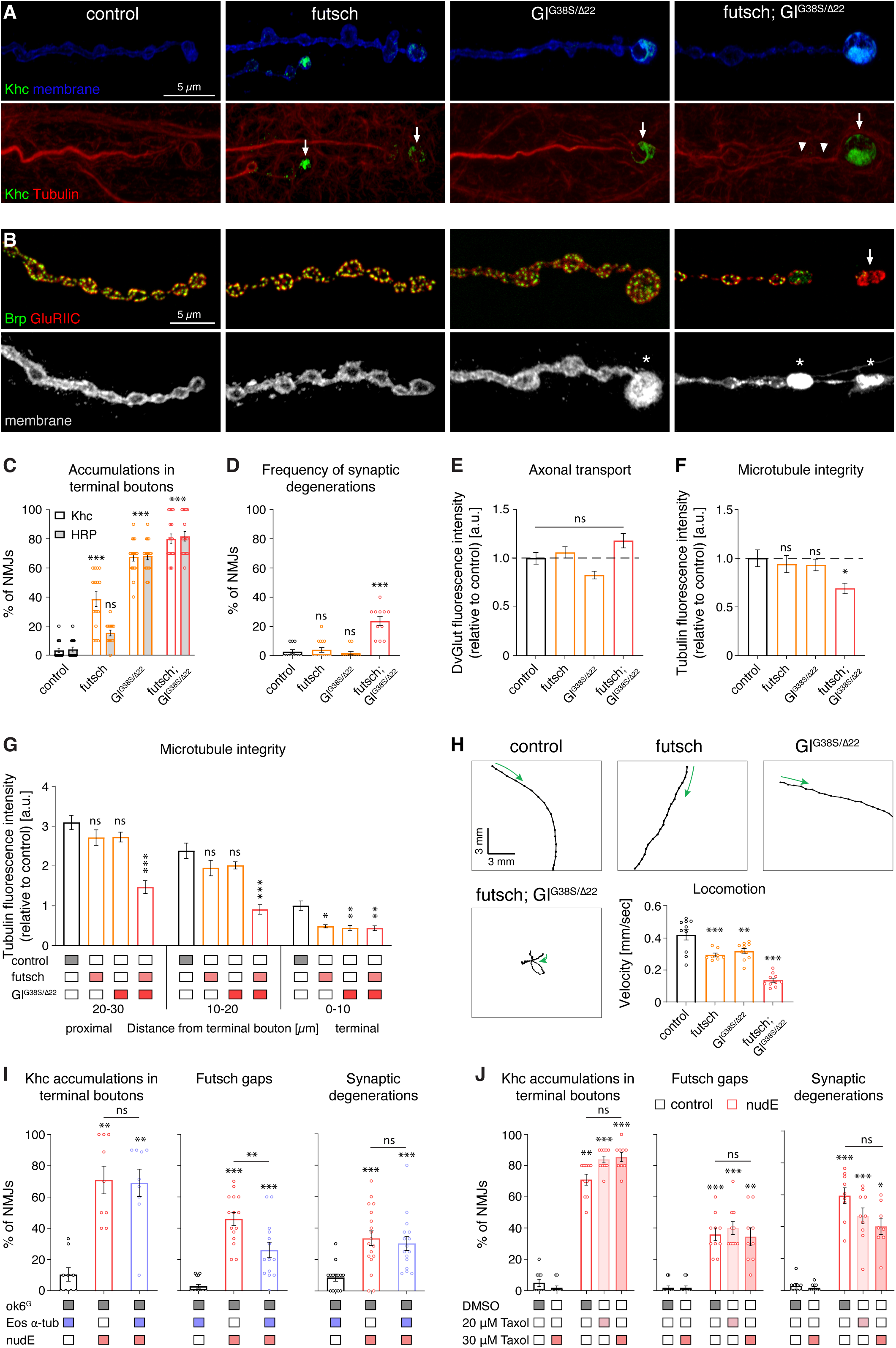
A two-hit model reveals reciprocal dependence between transport initiation and microtubule maintenance. (A) Confocal images of muscle 4 NMJs stained for Khc (green) and neuronal membrane (HRP, blue, top row) or Khc (green) and Tubulin (red, bottom row) in control, *futsch^K68^* single mutant, *Gl^G38S^*^/*Δ22*^ single mutant, and *futsch^K68^*; *Gl^G38S^*^/*Δ22*^ double mutant animals. *futsch^K68^* mutants show Khc accumulations at terminal boutons (arrows) without HRP accumulations. *Gl^G38S^*^/*Δ22*^ mutants show stronger Khc and HRP accumulations. Double mutants display enhanced accumulations and disrupted microtubule integrity (arrowheads). Scale bar, 5 µm. (B) Confocal images of the same genotypes stained for Brp (green), GluRIIC (red), and neuronal membrane (HRP, gray). Synaptic degeneration with membrane fragmentation (arrows) is only observed in double mutants. Scale bar, 5 µm. (C) Frequency of NMJs with Khc and HRP accumulations at terminal boutons. (D) Frequency of synaptic degeneration. Neither single mutant shows significant degeneration; the double mutant crosses the threshold. (E) Axonal transport measured by DvGlut fluorescence intensity in nerve bundles. No significant defects in any genotype. (F) Overall Tubulin fluorescence intensity at the NMJ. (G) Tubulin intensity within the last 30 µm of muscle 4 NMJs, measured in 10 µm intervals. Single mutants show reduction only at the distal-most 0–10 µm; the double mutant shows significant reduction across the entire terminal region. (H) Representative locomotion trajectories (left) and quantification of velocity (bottom). Green arrows indicate starting point and direction. Both single mutants show mild locomotion defects that are significantly enhanced in the double mutant. (I) Motoneuron expression of *Eos-α-tubulin* in *nudE* mutants (*ok6*-Gal4>UAS-*Eos-α-tubulin*; *nudE^39A^*^/*39A*^). Increasing the available Tubulin pool significantly reduces Futsch gaps (middle) but does not rescue Khc accumulations at terminal boutons (left), and synaptic degeneration shows only a marginal trend toward improvement (right). (J) Feeding of *nudE* mutants with the microtubule-stabilizing drug Taxol (20–30 µM). Neither concentration rescues Khc accumulations, Futsch gaps, or synaptic degeneration. (C-J) Mean ± s.e.m. two-way ANOVA with Sidak’s and Tukey’s multiple comparisons test (C, Sidak’s, G, Tukey’s); Kruskal-Wallis test with Dunn’s multiple comparisons test (D, I, J); one-way ANOVA with Sidak’s multiple comparisons test (E, F, H, I Futsch gaps). ns, p ≥ 0.05; *p < 0.05; **p < 0.01; ***p < 0.001. n = 15, 15, 17, 17 animals (C); n = 11, 15, 11, 11 animals (D); n = 64, 57, 65, 49 nerve bundles (E); n = 35, 35, 35, 30 NMJs (F); n = 28, 25, 29, 29 NMJs (G); n = 10, 8, 10, 10 animals (H); n = 9 animals per genotype (I, Khc), n = 15, 15, 14 animals (I, Futsch gaps); n = 14, 18, 16 animals (I, synaptic degenerations); n = 10,11, 10, 10, 9 (J, Khc accumulations), n = 11, 11, 10, 10, 9 (J, Futsch gaps), n = 9, 10, 9, 10, 8 (J, synaptic degenerations).

To further test the two-hit model, we combined each of the two mutations with a mild reduction of presynaptic Tubulin levels. Weak neuronal expression of a *β-tubulin* RNAi construct did not affect synaptic stability, axonal transport, or overall microtubule abundance (Figure S5A–F; D42-Gal4). However, all three single mutant conditions, *futsch^K68^*, *Gl^G38S^*, and *β-tub^RNAi^*, caused Khc accumulations at terminal boutons (38.7%, 67.7%, and 39.4% respectively; Figure 7C, S5C). The observation that even mild perturbations of the microtubule cytoskeleton (*futsch^K68^*and *β-tub^RNAi^*) result in transport initiation defects indicates that the precise levels of microtubules and microtubule-associated proteins at the synaptic terminal are critical for the normal initiation of retrograde transport. When *β-tub^RNAi^* was combined with either *futsch^K68^* or *Gl^G38S^*, both double mutant conditions crossed the threshold into significant synaptic degeneration (*futsch^K68^*; *β-tub^RNAi^*: 35.5±5.0%; *β-tub^RNAi^*; *Gl^G38S^*: 41.7±4.1%), with severe disruption of microtubule integrity and impaired locomotion (Figure S5A–J). Thus, three independent genetic combinations converge on the same outcome: sub-threshold perturbations that individually affect either retrograde transport initiation or the microtubule cytoskeleton can synergistically cause synaptic degeneration when combined. Our finding that transport initiation failure in *nudE* mutants leads to secondary microtubule loss (Figure 5) and our observation that microtubule perturbations themselves impair transport initiation together reveal a reciprocal dependence between retrograde transport initiation and microtubule maintenance at the synaptic terminal. A sufficiently strong defect in either process, as in *nudE* null mutants or in the dynein complex knockdown conditions, is predicted to eventually compromise the other, crossing the threshold into progressive degeneration.

The reciprocal dependence model predicts that increasing microtubule levels in *nudE* mutants should improve microtubule integrity but not rescue the primary transport initiation defect. To test this, we expressed Eos-tagged α-Tubulin in motoneurons of *nudE* mutant animals. Increasing the available Tubulin pool significantly reduced the frequency of Futsch gaps (from 45.8±4.1% to 26.0±4.9%) but did not improve Khc accumulations at terminal boutons (71.0±8.9% vs 69.2±8.7%), and synaptic degeneration showed only a marginal trend toward improvement (33.6±4.7% vs 30.3±4.4%) (Figure 7I). Feeding of *nudE* mutant larvae with the microtubule-stabilizing drug Taxol (20–30 µM) similarly failed to rescue any parameter (Figure 7J). Thus, while reducing microtubule levels impairs transport initiation (Figure 7C, S5C), increasing the free Tubulin pool does not have the inverse effect. This asymmetry indicates a precise regulation of microtubule quantity and organization at the synaptic terminal to balance structural support and transport initiation; excess Tubulin can be incorporated into additional polymer, but this does not compensate for the failure to activate the retrograde transport machinery. Moreover, Taxol-mediated stabilization suppresses the dynamic microtubule plus-ends on which the EB/CLIP-170/dynactin recruitment cascade depends (Moughamian et al., 2013), which would further impair rather than restore transport initiation. These results confirm that the failure to initiate retrograde transport is the primary defect in *nudE* mutant motoneurons, and that the resulting progressive loss of microtubule integrity and synaptic stability cannot be overcome by manipulating microtubule levels or stability alone.

## Discussion

Recent *in vitro* reconstitution studies established that the mammalian ortholog Nde1 functions as a transient scaffold that tethers Lis1 to autoinhibited dynein, enabling the formation of a structural intermediate that is rate-limiting for dynein activation (Singh et al., 2024; Yang et al., 2026; Zhao et al., 2023). However, whether NudE/Nde1 is required specifically for transport initiation in neurons had not been tested. Here, we demonstrate that the dynein regulator NudE is essential for retrograde transport initiation at the synaptic terminal, and that failure of this process triggers a progressive cascade of microtubule destabilization and synaptic degeneration. First, NudE loss causes accumulation of dynein and kinesin motor proteins at distal synaptic boutons while preserving transport velocity, the defining signature of a transport initiation defect (Lloyd et al., 2012; Moughamian and Holzbaur, 2012). Second, our temporal analysis reveals that transport initiation failure is the earliest detectable defect, preceding both synaptic degeneration and axonal transport disruption. Third, structure-function analysis demonstrates that the biochemically defined Dic-binding interface of NudE (Okada et al., 2023; Singh et al., 2024; Wang and Zheng, 2011; Zyłkiewicz et al., 2011) is required for transport initiation *in vivo*, with graded disruption of this interface producing correspondingly graded phenotypes. Fourth, perturbation of any dynein-dynactin complex component phenocopies NudE loss, consistent with the requirement for stoichiometric dynein motor complex assembly. Thus, our data provide the *in vivo* validation that NudE/Nde1 is required for axonal transport initiation and that the consequences of failed dynein activation extend far beyond a transport deficit, they initiate a degenerative cascade that recapitulates key features of dying-back neuropathies.

A remarkably similar degenerative phenotype has been observed in the mammalian system: postnatal deletion of the ortholog *Ndel1* in mouse forebrain excitatory neurons causes microtubule fragmentation in CA1 hippocampal dendrites, progressive loss of synaptic contacts, and a dramatically shortened lifespan (Gavrilovici et al., 2021; Jiang et al., 2016; Kiroski et al., 2020). Interestingly, a single injection of the glycoprotein Reelin into the hippocampus partially rescues microtubule integrity, dendritic morphology, and synapse number, and partially extends the lifespan of *Ndel1* conditional knock-out animals (Jiang et al., 2016; Kiroski et al., 2020). However, the interpretation of these findings is complicated by the fact that Reelin signals through the Dab1/GSK3β pathway to regulate microtubule stability independently of dynein (Jiang et al., 2016), leaving open whether the *Ndel1* phenotype reflects failed dynein activation, disrupted Reelin signaling, or both. Our analysis in Drosophila motoneurons resolves this ambiguity. Although a Reelin/F-spondin-related protein, Drospondin, has recently been identified in Drosophila glia (Rojo-Cortés et al., 2026), the canonical Reelin-Dab1 signaling cascade is absent. The phenotypic convergence between Ndel1 loss in mouse hippocampal neurons and NudE loss in Drosophila motoneurons therefore reflects the conserved dynein activation function rather than shared downstream signaling. The partial rescue by Reelin in the mouse is consistent with Reelin acting on microtubule stability via the Dab1/GSK3β pathway, analogous to the partial rescue we observe upon *tubulin* overexpression in *nudE* mutants: both interventions address the secondary microtubule defect without restoring dynein-dependent transport initiation.

A central finding of this study is the reciprocal dependence between retrograde transport initiation and microtubule maintenance at the synaptic terminal. Our temporal and genetic analyses demonstrate that transport initiation failure precedes microtubule destabilization, establishing the directionality of the cascade. However, the two-hit experiments reveal that the relationship is not unidirectional: even mild perturbations of the microtubule cytoskeleton, through loss of the microtubule-associated protein Futsch/MAP1B or reduction of Tubulin levels, are sufficient to impair retrograde transport initiation. This bidirectional dependence creates a vulnerability in which a sub-threshold defect in either process can be amplified by its effect on the other, ultimately crossing the threshold into synaptic degeneration. Consistent with this model, three independent genetic combinations that individually remain below the degeneration threshold synergistically cause synaptic loss when combined. The dependence of transport initiation on microtubule integrity is consistent with the finding that dynactin is recruited to the distal axon through an ordered cascade that requires dynamic microtubule plus-ends: EB proteins and CLIP-170 capture growing microtubules and recruit dynactin via its CAP-Gly domain, while Lis1 independently activates the dynein motor (Moughamian et al., 2013; Moughamian and Holzbaur, 2012). NudE, by tethering Lis1 to dynein (Yang et al., 2026; Zhao et al., 2023), operates in the motor activation arm of this pathway. In this framework, perturbations that reduce microtubule dynamics at the terminal, whether through loss of MAPs, reduced Tubulin availability, or progressive lattice damage, would diminish the substrate on which the EB/CLIP-170/dynactin recruitment cascade depends, thereby impairing transport initiation even when NudE-Lis1-dynein function is intact. A sufficiently severe defect in either arm, as in *nudE* null mutants, is predicted to compromise the other, explaining why complete loss of NudE function produces both transport initiation failure and secondary microtubule destabilization within a single genetic perturbation.

An unresolved question is why transport initiation failure leads to progressive microtubule loss rather than simply causing cargo accumulation. Recent *in vitro* studies have revealed that molecular motors, including single dynein molecules, damage the microtubule lattice as they walk, removing Tubulin dimers from the shaft (Triclin et al., 2021). Free GTP-Tubulin dimers incorporate into these damaged sites, creating rescue sites that protect microtubules from catastrophe and increase their lifespan (Andreu-Carbó et al., 2022; Triclin et al., 2021). Our mitochondrial transport data are consistent with this hypothesis: in *nudE* mutants, both the proportion of retrogradely moving mitochondria and their velocity are reduced, while total mitochondrial density remains unchanged (Figure 2I, J, S2C). The resulting decrease in cumulative retrograde motor traffic would reduce the frequency of motor-induced repair events along synaptic microtubules. A similar cargo-specific requirement for NDEL1 in retrograde mitochondrial transport has been reported in mammalian sensory neurons (Pandey et al., 2022). This motor-induced repair generates a positive feedback loop: microtubules that carry more motor traffic accumulate more rescue sites and become selectively stabilized (Andreu-Carbó et al., 2022). Tubulin incorporation into the lattice of pre-existing microtubules has recently been confirmed to occur in living cells (Gazzola et al., 2023), and mechanical forces including compressive stress can further stabilize microtubules through the recruitment of lattice-protective factors such as CLASP2 (Li et al., 2023). Although this repair mechanism has not yet been examined in neurons, it raises the possibility that a similar process operates at the synaptic terminal. In this scenario, the absence of NudE-dependent dynein activation would severely diminish retrograde motor transport along synaptic microtubules, reducing the motor-induced repair that normally maintains microtubule integrity. Without ongoing repair, lattice damage would accumulate progressively, leading to the distal-to-proximal gradient of microtubule loss and the eventual synaptic degeneration that we observe. This model would also explain why artificially stabilizing microtubules with Taxol or increasing the free Tubulin pool fails to rescue the transport initiation defect, the problem is not insufficient microtubules but the absence of the dynamic motor-microtubule interaction that maintains lattice integrity over time.

The progressive, length-dependent synaptic degeneration in *nudE* mutants shares key features with dying-back neuropathies in humans, in which distal nerve terminals degenerate first and the disease progresses proximally over time (Moloney et al., 2014; Schaumburg et al., 1974). A major unresolved question in these diseases is whether axonal transport defects are a cause or a consequence of degeneration (Berth and Lloyd, 2023). Genetic evidence from mouse models strongly supports a causal role since disruption of the dynein-dynactin complex by *dynamitin* overexpression in postnatal motor neurons causes late-onset progressive motor neuron degeneration (LaMonte et al., 2002) and missense mutations in the dynein heavy chain produce a similar phenotype (Hafezparast et al., 2003). Our temporal analysis extends this evidence by resolving the sequence of events at the level of individual synapses: transport initiation failure is detectable before structural degeneration becomes apparent, establishing it as an initiating event. The distal synapse is likely the most vulnerable site because this is where retrograde cargo first engages the dynein motor complex; along the axon, transport can be reinitiated at multiple points, providing greater resilience.

The threshold behavior we observe, where sub-threshold transport initiation defects are tolerated until a second perturbation tips the balance, may explain why many neurodegenerative diseases manifest only after years of sub-threshold functional impairment. The *Drosophila* HMN7B model provides a striking example: the *Gl^G38S^* mutation causes transport initiation defects at larval synapses without triggering degeneration, yet adult animals develop progressive motor deficits and early lethality (Lloyd et al., 2012). Similarly, postnatal ablation of the full-length *p150^Glued^*protein in mouse neurons causes no developmental phenotype, but 12 months later, it leads to selective spinal motor neuron degeneration (Yu et al., 2018). These observations suggest that chronic, sub-threshold transport initiation defects gradually compromise microtubule integrity and synaptic stability, progressively reducing the capacity for sustained synaptic transmission until structural degeneration ensues. The electrophysiological analysis at segment A3 provides direct evidence for this reduced capacity: baseline synaptic transmission is normal, but repetitive stimulation at 10 Hz, well below the endogenous motoneuron burst frequency of 20–30 Hz during larval locomotion (Ormerod et al., 2022), reveals rapid rundown and frequent transmission failures, indicating that the functional integrity of the synapse is already compromised before structural degeneration becomes apparent. Motor neurons, with their exceptionally long axons, would be the first to cross this threshold, consistent with the preferential vulnerability of motor neurons in both the p150Glued mouse model (Yu et al., 2018) and in human diseases caused by *dynein-dynactin* mutations (Farrer et al., 2009; Puls et al., 2003).

Several aspects of the proposed molecular mechanism underlying dying-back neurodegeneration in our model remain to be resolved. While we establish that transport initiation failure precedes microtubule loss, the suggested repair-based molecular mechanism linking reduced motor traffic to microtubule destabilization in neurons has yet to be experimentally validated, and we cannot exclude that the accumulation of cargo and membrane at terminal boutons contributes to microtubule damage through mechanical stress. Testing whether motor-induced lattice repair (Andreu-Carbó et al., 2022; Triclin et al., 2021) operates at synaptic terminals will be an important future goal. We also note that impaired retrograde transport will inevitably affect anterograde transport, as kinesin motors depend on dynein-mediated recycling, and disruption of anterograde transport can itself cause synaptic retractions (Valakh et al., 2012). However, our two-hit conditions produce synaptic degeneration without significant axonal transport defects, indicating that degeneration can arise from the combined perturbation of retrograde transport initiation and microtubule integrity before general axonal transport is detectably impaired.

In summary, our work identifies NudE as a critical regulator of retrograde transport initiation at the synaptic terminal and establishes a mechanistic link between failed dynein activation and progressive neurodegeneration. The cascade we describe, transport initiation failure followed by microtubule destabilization, impaired synaptic transmission, and structural degeneration, provides a framework for understanding how defects in a single molecular pathway can produce the progressive, length-dependent degeneration observed in dying-back neuropathies. The reciprocal dependence between transport initiation and microtubule maintenance, and the threshold behavior that governs the gradual transition from functional impairment to structural degeneration may have broad implications for understanding the onset and progression of neurodegenerative diseases associated with dynein-dynactin dysfunction.

## Acknowledgments

We thank the Bloomington Drosophila Stock Center (NIH P40OD018537) for fly stocks, the Developmental Studies Hybridoma Bank (DSHB, University of Iowa) for antibodies, and the BACPAC Resources Center (Children’s Hospital Oakland Research Institute) for P[acman] BAC clones. We are grateful to the Hays lab (University of Minnesota) for providing the Dhc antibody, and to the Kolodkin lab and Thomas Lloyd for providing Glued fly stocks. We thank all members of the Pielage lab for helpful discussions. This work was supported by the Deutsche Forschungsgemeinschaft (DFG; GRK2737 STRESSistance) (T.M. and J.P.) and the Forschungsinitiative Rheinland-Pfalz BioComp (T.M. and J.P.).

## Author contributions

Z.M., R.S. and J.P. conceived and designed the study. Z.M. performed all experiments and analyzed all data with assistance from D.A.L., L.M.L. and J.E.; B.E. performed the *in vivo* live-imaging experiments. R.S. and D.B. performed initial experiments and generated *nudE* constructs with E.M.; T.M. provided critical input. Z.M. and J.P. wrote the manuscript with input from all authors.

## Data availability

All data and genetic tools are available from the corresponding author.

## Methods

### Fly stocks

*Drosophila melanogaster* stocks used in this study are listed in Table 1. Stocks were obtained from the Bloomington Drosophila Stock Center (BDSC) and the Vienna Drosophila Resource Center (VDRC) unless otherwise indicated. All stocks were raised on standard Cold Spring Harbor Protocols (CSHL) fly food at room temperature, and genetic crosses were maintained at 25°C with 65±5% humidity unless otherwise mentioned in figure legends. For control genotypes, *w*^1118^ was crossed to the respective parental genotype unless otherwise indicated in figure legends.

**Table 1.** Fly stocks used in this study.

| Genotype | Source | Identifier | Figure(s) |
| --- | --- | --- | --- |
| *w*<sup>1118</sup> | BDSC | #5905 | 1–7, S1–S5 |
| *nudE*<sup>39A</sup> | (Wainman et al., 2009) | N/A | 1–7, S1–S5 |
| Df(3L)BSC282 | BDSC | #23667 | S1 |
| BAC-*nudE* (short) | This study | N/A | S1 |
| BAC-*nudE* (long) | This study | N/A | S1 |
| *ok6*-Gal4 | BDSC | #64199 | 1, 2, 4–7, S2, S4 |
| *mef2*-Gal4 | BDSC | #27390 | 1, 6 |
| *elav*<sup>C155</sup>-Gal4 | BDSC | #458 | 1, 2, S3 |
| 5053A-Gal4 | BDSC | #2702 | 2 |
| *da*-Gal4 | BDSC | #55851 | S1D |
| *D42*-Gal4 | BDSC | #8816 | S5 |
| *UAS-dicer2* | BDSC | #24650 | S3, S5 |
| *UAS-nudE* (WT) | This study | N/A | 1–3, 5, 6, S1, S2, S4 |
| *UAS-nudE*<sup>E34A/E37A</sup> (Dic2) | This study | N/A | 3 |
| *UAS-nudE*<sup>E46A/E48A/D50A</sup> (Dic3) | This study | N/A | 3 |
| *UAS-nudE*<sup>ΔDic</sup> (Dic2+Dic3) | This study | N/A | 3 |
| *UAS-nudE*<sup>ΔLis1</sup> | This study | N/A | 3 |
| UAS-*nudE*<sup>ΔDhc</sup> | This study | N/A | 3 |
| 10xUAS-IVS-mCD8-GFP | BDSC | #32186 | 4 |
| UAS-Brp-mStraw | (Owald et al., 2010) | N/A | 4 |
| UAS-mitoGFP | BDSC | #8443 | 2, S2 |
| UAS-Eos-α-tubulin | BDSC | #51313 | 7 |
| UAS-*Dic*<sup>RNAi</sup> | VDRC | #v48334 | S3 |
| UAS-*Dlic*<sup>RNAi</sup> | VDRC | #v41686 | S3 |
| UAS-*Dhc*<sup>RNAi</sup> | VDRC | #v28054 | S3 |
| UAS-*Dmn*<sup>RNAi</sup> | VDRC | #v110741 | S3 |
| UAS-*Lis1*<sup>RNAi</sup> | VDRC | #v106777 | S3 |
| UAS-*Gl*<sup>DN</sup> | BDSC | #51645 | S3 |
| UAS-β-*tub*<sup>RNAi</sup> | VDRC | #v24138 | S5 |
| *futsch*<sup>K68</sup> | (Hummel et al., 2000) | N/A | 7, S5 |
| *Gl*<sup>G38S</sup> | (Lloyd et al., 2012) | N/A | 7, S5 |
| *Gl*<sup>Δ22</sup> | (Lloyd et al., 2012) | N/A | 7, S5 |

### Immunohistochemistry

*Drosophila* larvae at second, early third, and third instar stages were dissected in pre-chilled standard HL3 saline and fixed for 5 minutes with Bouin’s fixative (Art# 6482.3, Carl Roth). Larvae were stained overnight with primary antibodies at 4°C, followed by washing in PBST containing 0.1% Triton X-100. Secondary antibodies were applied for 2 h at room temperature, followed by washing in PBST (0.1% Triton X-100). Larval preparations were mounted on glass slides in ProLong Gold antifade medium (P36934, Thermo Fisher). The following primary antibodies were used: mouse anti-Brp (nc82) 1:250 (Developmental Studies Hybridoma Bank [DSHB], Iowa), rabbit anti-dGluRIIC 1:3,000 (Pielage et al., 2011), mouse anti-Futsch (22C10) 1:500 (DSHB), rabbit anti-Futsch 1:1,000 (this study), rabbit anti-DvGlut 1:10,000 (Bulat et al., 2014), rabbit anti-Khc 1:300 (Cytoskeleton, AKIN01), guinea pig anti-DLG 1:700 (this study), mouse anti-Dhc 1:1,000 (McGrail and Hays, 1997), and mouse anti-acetylated Tubulin 1:1,000 (Sigma, T6793). HRP conjugates were used as follows: HRP–Alexa Fluor 488 and HRP–Alexa Fluor 568 (or Cy3) at 1:1,000, and HRP–Alexa Fluor 647 at 1:500 (all Jackson ImmunoResearch Laboratories). Alexa Fluor 488/568-conjugated secondary antibodies were used at 1:1,000 and Alexa Fluor 647-conjugated secondary antibodies at 1:500 (Invitrogen). Images were acquired using an LSM700 confocal microscope (Zeiss) with 40×/1.3 NA and 63×/1.4 NA oil immersion objectives, and a Stellaris 8 confocal microscope (Leica) with a 20×/0.75 NA glycerol immersion and 63×/1.4 NA oil immersion objective. All genotypes within a single experiment were stained together and imaged under identical settings. For intensity quantifications, acquisition parameters were set using the respective control genotype and applied to all other genotypes. Image processing and analysis were performed using Imaris (Bitplane), Fiji (ImageJ), and Photoshop (Adobe); only linear adjustments were applied.

### Axonal transport live imaging

Axonal transport of mitochondria was monitored in motoneuron axon bundles of third instar larvae at segments A3/A4. Wandering third instar larvae were dissected on a Sylgard block in HL3. The Sylgard block with the dissected larva was inverted onto a glass-bottom 35 mm µ-Dish (Ibidi) containing HL3 supplemented with 100 µM 1-naphthylacetyl spermine trihydrochloride (NASPM; Sigma-Aldrich) to prevent muscle contractions. The Sylgard block was held in place with magnetic pins. Anterograde and retrograde transport of mitochondria was monitored by time-lapse imaging on a Zeiss LSM700/710 confocal microscope with a 63x/1.4 NA oil immersion objective.

Confocal image series were converted to movies using Fiji (ImageJ). Selected axons were stabilized with Fiji plugin MultiStackReg (Transformation: Translation and if necessary Scaled Rotation)(https://github.com/miura/MultiStackRegistration/blob/6dbf10620a8d8163742aa9abad6d f3524afa4bfc/src/main/java/de/embl/cmci/registration/MultiStackReg_.java). Kymographs, manual transport tracking and automated measurement of traced transport was done using the Fiji Plugin Kymolyzer (2020) (https://currentprotocols.onlinelibrary.wiley.com/doi/10.1002/cpcb.107) (https://github.com/ThomasSchwarzLab/KymolyzerCodes).

### Electrophysiological recordings

Sharp electrode intracellular recordings were performed in bridge mode on third instar larvae as described previously (Choudhury et al., 2016). Briefly, wandering third instar larvae were dissected in pre-chilled Ca^2+^-free HL3 (70 mM NaCl, 5 mM KCl, 20 mM MgCl₂, 10 mM NaHCO₃, 115 mM sucrose, 5 mM trehalose, 5 mM HEPES, 2 mM EGTA) on 35 mm Sylgard plates. Recordings were performed in HL3 containing 0.5 mM Ca^2+^ and 0 mM EGTA using sharp electrodes with a resistance of 15–20 MΩ. Miniature excitatory junctional potentials (mEJPs) were collected for 60 s, and at least 30 evoked excitatory junctional potentials (EJPs) were recorded by stimulating the segmental nerves of the respective hemisegments with suprathreshold pulses delivered by an isolated pulse stimulator (A-M Systems) at 1 Hz. Only recordings with a muscle input resistance ≥ 3 MΩ and a resting membrane potential ≤ −60 mV were included in the analysis. Following single-stimulus recordings, the preparation was rested for one minute without stimulation. High-frequency stimulation (HFS) was then delivered at 10 Hz for 300 s. Only recordings maintaining a stable baseline (maximum shift of ±5%) were included. Recordings were performed on muscle 6/7 and muscle 4 NMJs of segments A3 and A6. Signals were amplified using an AxoClamp 900A (Molecular Devices) and digitized with a Digidata 1550B (Molecular Devices).

mEJP and EJP amplitudes were analyzed using MiniAnalysis software (Synaptosoft). Both mEJPs and EJPs were averaged per muscle cell. Quantal content was calculated by dividing the mean EJP amplitude by the mean mEJP amplitude for each cell. EJP amplitudes during high-frequency stimulation were quantified manually using the Threshold Search function in Clampfit 10.7 (Molecular Devices). EJP amplitudes were sampled every 30 s and averaged across three consecutive events at each time point.

### Generation of nudE constructs

The full-length wild-type *nudE* open reading frame (ORF) was amplified from cDNA LD19982 obtained from the *Drosophila* Genomics Resource Center (Indiana, USA) and was cloned into pENTR-D-TOPO using Gateway cloning and exchanged into pUASattB-10xUAS destination vectors using LR clonase (Invitrogen). Point mutations disrupting binding to the dynein intermediate chain (Dic) and deletions of the Lis1-binding and dynein heavy chain (Dhc)-binding domains were introduced into the wild-type *nudE* ORF using the QuikChange II site-directed mutagenesis kit (Agilent Technologies) following standard protocols. All constructs were exchanged into pUASattB-10xUAS destination vectors using LR clonase, sequence-verified, and injected into the *Drosophila* genome by standard φC31-mediated transgenesis into *attP40* to ensure equal expression levels (BestGene Inc., California, USA). Primers used to generate all constructs are listed in Table 2. The genomic P[acman] rescue constructs spanning the nudE genomic locus (CH321-75F7; CH322-121J07) were obtained from the P[acman] BAC libraries (BACPAC Resources Center; (Venken et al., 2009)) and integrated into the *attP40* site by standard φC31-mediated transgenesis (BestGene Inc., California, USA).

**Table 2.**
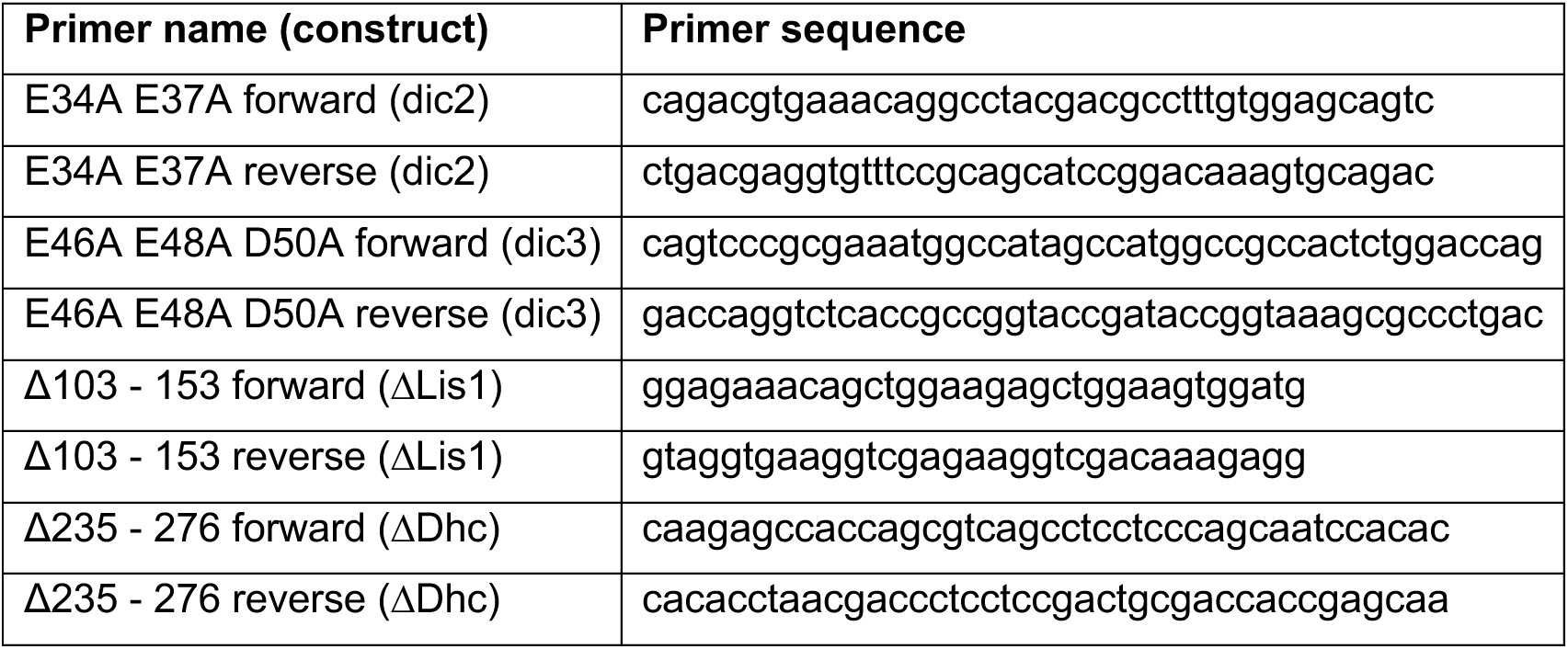
Primers for generation of *nudE* point mutations and deletions.

### Live non-invasive imaging

To non-invasively monitor synaptic degeneration throughout larval development, larvae were anesthetized in a 35 mm Ibidi µ-Dish using 30 µl of desflurane (Suprane, Baxter). Anesthetized larvae were mounted in 10S Voltalef halocarbon oil (VWR) under a glass coverslip. Images were acquired using a spinning-disc confocal microscope (Olympus IX81) with a 40× oil immersion objective. Muscle 26 and 27 NMJs were selected for imaging because they lie between cuticle and muscle and are readily accessible to optical microscopy. Larvae were returned to fly food supplemented with yeast between imaging sessions and exposed to laser illumination for a maximum of 15 minutes per session. Images were deconvolved using Huygens Remote Manager (Scientific Volume Imaging B.V.), aligned, stitched, and converted to maximum intensity projections using custom macros for Fiji (ImageJ).

### Larval crawling

Larval locomotion was assessed using an established crawling assay (Cunningham et al., 2018; Perry et al., 2017). Wandering third instar larvae were collected, rinsed in PBS, and allowed to acclimate on 3.5% solidified agarose plates for 1–2 minutes. Larvae were then transferred to a fresh agarose plate and recorded using a digital camera (Canon) at 1920 × 1088-pixel resolution and 25 fps for 3 minutes or until the larva reached the edge of the plate. Each larva was tested for three trials, and the averaged data were represented as a single data point.

### Taxol treatment

Taxol (paclitaxel) was prepared as a 10 mM stock solution in DMSO. Freshly prepared food vials were thoroughly mixed with Taxol and allowed to stand at room temperature for 12 h. Second instar larvae were collected from standard fly food and transferred to food containing 20 or 30 µM Taxol and 0.1% DMSO. Vials containing only 0.1% DMSO served as vehicle controls. Larvae were reared on Taxol-containing food at 25°C for 72 h before dissection. To ensure that only fed larvae were used for analysis, Taxol and DMSO-containing food was supplemented with blue food coloring dye.

### Quantification of phenotypes

#### Synaptic degeneration

Synaptic degeneration was quantified using an established synaptic degeneration assay (Eaton et al., 2002; Mushtaq et al., 2022; Pielage et al., 2005). Synaptic degenerations were identified by the presence of the postsynaptic glutamate receptor marker GluRIIC in the absence of the presynaptic active zone marker Brp. The frequency of synaptic degeneration represents the percentage of NMJs with degeneration per animal. The severity of synaptic degeneration was classified by the number of postsynaptic bouton profiles lacking presynaptic Brp: 1–2 boutons (score 1), 3–6 boutons (score 2), ≥7 boutons or total elimination (score 3). Quantifications were performed on muscles 1/9, 2/10, 4, and 6/7 of hemisegments as indicated in the figure legends and supplementary data table.

#### Axonal transport

Axonal transport defects were quantified in two independent ways. First, DvGlut accumulations in axon bundles stained with HRP and DvGlut were quantified under a TCS SPE DM5500Q microscope (Leica) with a 40×/0.75 NA air objective. Second, axon bundles from larvae stained together with HRP, DvGlut, and Brp were imaged under identical settings using a Leica Stellaris 8 microscope and 63x/1.4 NA oil immersion objective. The integrated density of DvGlut and Brp was measured using a custom macro for Fiji (ImageJ). HRP signal was used to mask axon bundle area from background.

#### Retrograde transport initiation

Defects in retrograde transport initiation were quantified as accumulation of the anterograde motor protein Kinesin heavy chain (Khc) and HRP at boutons (predominantly terminal boutons) of NMJs labeled with Khc, HRP, and DLG. In control animals, neither Khc nor Dhc were detectable by confocal microscopy at synaptic boutons, reflecting an even distribution of motor complexes at low abundance and their rapid turnover by retrograde transport. Data represent the percentage of NMJs with Khc/HRP accumulations at terminal boutons per animal.

#### Futsch gaps

Futsch gaps were quantified under a TCS SPE DM5500Q microscope (Leica) with a 40×/0.75 NA air objective on NMJs labeled with Futsch and HRP. Any discontinuity in the Futsch structure within the HRP-labeled neuronal membrane was scored as a Futsch gap.

#### Fluorescence intensity measurements

Fluorescence intensity of individual proteins at axon bundles or NMJs was measured on maximum intensity projections using a custom macro for Fiji (ImageJ). All larvae within an experiment were stained together and imaged under identical settings. Acquisition parameters were set using the respective control genotype and applied to all other genotypes. HRP signal was used to delineate neuronal membrane at axon bundles or NMJs, and background fluorescence was subtracted from the signal in each image. The integrated density of Tubulin within the last 30 µm of the NMJ (Figure 5H, I; Figure 7G; Figure S5G) was measured manually on maximum intensity projections using Fiji (ImageJ).

#### Futsch and Tubulin intensity at Futsch gaps

Tubulin was not completely absent at Futsch gaps in *nudE* mutant NMJs. To determine Tubulin levels at sites with and without Futsch gaps, maximum intensity projections of NMJs stained for Futsch, acetylated-Tubulin, and HRP were analyzed using the Plot Profile tool in Fiji (ImageJ). A line encompassing the diameter of the NMJ, defined by HRP, was drawn to generate intensity profiles for Futsch and acetylated-Tubulin. For control and rescue animals, profile plots were acquired at a minimum of five regularly spaced positions per NMJ. For *nudE* mutants, profiles were acquired at Futsch gap sites, and adjacent Futsch-positive regions served as internal controls. The number of profiles per *nudE* NMJ varied depending on the number of detectable gaps.

#### Larval crawling

Distance traveled was quantified manually from high-resolution videos (FFMPEG format) using the MTrackJ plugin in Fiji (ImageJ). Head turns were counted manually; only turns exceeding 70° were included. Each larva was tested for three trials, and the averaged data were represented as a single data point.

### Statistical analysis

Statistical analyses were performed using GraphPad Prism 11. All data were tested for normal distribution using D’Agostino–Pearson omnibus and Shapiro–Wilk normality tests. Parametric (one-way or two-way ANOVA) or non-parametric tests were applied accordingly, with relevant post hoc corrections as indicated in the figure legends and supplementary data table. All statistical tests were two-tailed. F values, degrees of freedom, and exact p values for all comparisons are provided in the supplementary data table. Significance levels were defined as: ***p ≤ 0.001, **p ≤ 0.01, *p ≤ 0.05, and ns (non-significant) p ≥ 0.05.

