## Supplemental Data for "Loss of NudE-mediated dynein activation at synaptic terminals causes progressive axon length-dependent neurodegeneration"

#### Supplementary Figure Legends

**Figure S1: Additional characterization of synaptic degeneration in *nudE* mutants.** (A, B) Frequency (A) and severity (B) of synaptic degeneration on muscles 1/9, 2/10, 4, and 6/7 NMJs in *nudE* mutants (*nudE*$^{39A/39A}$ and *nudE*$^{39A/Df}$) compared to controls. All degeneration phenotypes are rescued by bacterial artificial chromosomes (BAC) encompassing the *nudE* locus. (C, E) Frequency (C) and severity (E) of synaptic degeneration along the anterior-posterior body axis on muscles 1/9, 2/10, 4, and 6/7 NMJs. (D) Table showing viability and larval locomotion of different *nudE* genotypes. (A-C, E) Mean ± s.e.m. Kruskal-Wallis test with Dunn's multiple comparisons test (A); two-way ANOVA with Tukey's multiple comparisons test (C, E). ns, $p \geq 0.05$; *$p < 0.05$; **$p < 0.01$; ***$p < 0.001$. n = 20, 20, 15, 12, 12 animals (A-E).

**Figure S2: Motor accumulations along the body axis and mitochondrial transport parameters**. (A) Confocal images of muscle 4 NMJs stained for Dynein heavy chain (Dhc, red), Kinesin heavy chain (Khc, green), and neuronal membrane (HRP, blue or gray). In controls, neither motor protein accumulates at terminal boutons. In *nudE* mutants (*nudE*$^{39A/39A}$), both Dhc and Khc accumulate at distal boutons alongside membrane-associated proteins. Scale bar, 5 µm. (B) Frequency of Khc (top) and HRP (bottom) accumulations at terminal boutons along the anterior-posterior body axis. All phenotypes are rescued by motoneuron expression of *nudE* (*nudE*, *ok6*-Gal4>UAS-*nudE*). (C) Mitochondrial density (number of mitochondria per 100 µm axon length) is not significantly different between control, *nudE* mutant, and *nudE* rescue (*nudE*, *5053A*-Gal4>UAS-*nudE*) animals. (D) Anterograde velocity of motile mitochondria is not significantly affected by loss of NudE. (E) Average anterograde travel length of motile mitochondria. (F) Average retrograde travel length of motile mitochondria. (B-F) Mean ± s.e.m. Two-way ANOVA with Tukey's multiple comparisons test (B, red stars indicate comparisons to the A2 data); one-way ANOVA with Sidak's multiple comparisons test (C-E); Kruskal-Wallis test with Dunn's multiple comparisons test (F). ns, $p \geq 0.05$; *$p < 0.05$; **$p < 0.01$; ***$p < 0.001$. n = 14, 14, 10 animals (B); n = 10, 10, 8 axons (C), n = 10, 7, 8 axons (D, E), n = 10, 9, 8 axons (F) from a minimum of 6 animals per genotype.

**Figure S3: Perturbation of dynein-dynactin complex components phenocopies *nudE* loss.** (A) Schematic of core dynein-dynactin complex subunits. (B–H) Confocal images of muscle 4 NMJs stained for Kinesin heavy chain (Khc, green) and neuronal membrane (HRP, red). Control (B) and motoneuron-specific knockdown of *Dic* (C), *Dlic* (D), *Dhc* (E), *Dmn/Dynamitin* (F), or dominant-negative expression of *p150*$^{Glued}$ (G), or *Lis1* knockdown (H). All perturbations cause Khc and membrane accumulations at terminal boutons. Scale bar, 5 µm. (I) Frequency of Khc and HRP accumulations at terminal boutons for each genotype. (J) Frequency of synaptic degeneration. In all genotypes, the frequency of Khc accumulations exceeds the frequency of degeneration. (K–Q)

Confocal images of nerve bundles stained for the active zone protein Brp (green), synaptic vesicle transporter DvGlut (red), and neuronal membrane (HRP, blue). Control (K) and motoneuron-specific knockdown of *Dic* (L), *Dlic* (M), *Dhc* (N), *Dmn/Dynamitin* (O), or dominant-negative expression of $p150^{Glued}$ (P), or *Lis1* knockdown (Q). Scale bar, 10 µm. (R) Fluorescence intensity of Brp and DvGlut in nerve bundles, normalized to controls. All larvae were reared at 28°C. (I, J, R) Mean ± s.e.m. Two-way ANOVA with Tukey's multiple comparisons test (I, R); One-Way ANOVA with Sidak's multiple comparisons test (J). ns, $p \geq 0.05$; *$p < 0.05$; **$p < 0.01$; ***$p < 0.001$. n = 10, 9, 10, 10, 10, 7, 10 animals (I); n = 10, 9, 9, 10, 10, 10, 10 animals (J); n = 41, 35, 46, 37, 37, 47, 49 nerve bundles from 8–12 animals per genotype (R).

**Figure S4: Raw electrophysiological values at segments A3 and A6.** Quantification of mEJP amplitude (A, D), EJP amplitude (B, E), and quantal content (C, F) at muscle 6/7 NMJs in proximal segment A3 (A–C) and distal segment A6 (D–F). In *nudE* mutants ($nudE^{39A/39A}$), synaptic transmission is significantly reduced at A6 but not at A3. All parameters are rescued by motoneuron expression of *nudE* (*nudE*, *ok6*-Gal4>UAS-*nudE*). (G–I) Quantification of mEJP amplitude (G), EJP amplitude (H), and quantal content (I) at muscle 4 NMJs in proximal segment A3. (J–L) Quantification of mEJP amplitude (J), EJP amplitude (K), and quantal content (L) at muscle 4 NMJs in distal segment A6. In *nudE* mutants, synaptic transmission is significantly reduced at A6 but not at A3 on muscle 4. (A-L) Mean ± s.e.m. One-way ANOVA with Sidak's multiple comparisons test (A-I, L); Kruskal-Wallis test with Dunn's multiple comparisons test (J, K). ns, $p \geq 0.05$; *$p < 0.05$; **$p < 0.01$; ***$p < 0.001$. n = 11, 15, 12 animals (A–F); n = 20, 12, 15 animals (G); n = 13, 12, 14 animals (H-I); n = 20, 11, 14 animals (J); n = 17, 11, 14 animals (K-L).

**Figure S5: Additional two-hit combinations confirm reciprocal dependence between transport initiation and microtubule maintenance**. (A) Confocal images of muscle 4 NMJs stained for Khc (green) and neuronal membrane (HRP, blue, top) or Khc (green) and Tubulin (red, bottom) in control, $\beta\text{-}tub^{RNAi}$, $futsch^{K68}$; $\beta\text{-}tub^{RNAi}$, and $\beta\text{-}tub^{RNAi}$; $Gl^{G38S/\Delta22}$ animals. $\beta\text{-}tub^{RNAi}$ alone causes mild Khc accumulations (arrows) without affecting microtubule integrity. Both double mutant combinations show enhanced Khc accumulations and a severe reduction of terminal microtubules (arrowheads). Scale bar, 5 µm. (B) Confocal images of the same genotypes stained for Brp (green), GluRIIC (red), and neuronal membrane (HRP, gray). Synaptic degeneration with membrane fragmentation (asterisks) is observed only in double mutant animals. Note that HRP accumulations are present after $\beta\text{-}tub^{RNAi}$ expression, but the membrane remains continuous. Scale bar, 5 µm. (C) Frequency of Khc accumulations at terminal boutons. (D) Frequency of synaptic degeneration. (E) Axonal transport measured by DvGlut fluorescence intensity. (F) Tubulin fluorescence intensity at the NMJ. (G) Tubulin intensity within the last 30 µm of muscle 4 NMJs, measured in 10 µm intervals. (H) Volume-rendered confocal image of a control muscle 4 NMJ labeled for Futsch (red) and

Tubulin (green), with white boxes indicating quantified areas. (I) Representative locomotion trajectories. Green arrows indicate starting point and direction. (J) Crawling velocity. (C-G, J) Mean ± s.e.m. Kruskal-Wallis test with Dunn's multiple comparisons test (C, E, F, J); One-way ANOVA with Sidak’s multiple comparisons test (D); two-way ANOVA with Tukey’s multiple comparisons test (G). ns, $p \geq 0.05$; *$p < 0.05$; **$p < 0.01$; ***$p < 0.001$. n = 15, 16, 17, 17 animals (C); n = 11, 15, 13, 9 animals (D); n = 64, 56, 49, 47 nerve bundles (E); n = 35, 33, 28, 33 NMJs (F); n = 28, 25, 27, 33 NMJs (G); n = 10 animals per genotype (J).

### Supplemental Figure S1

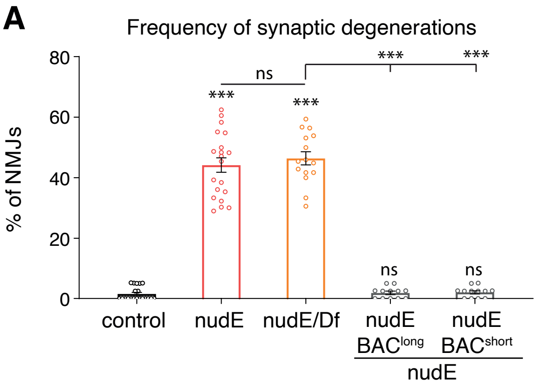


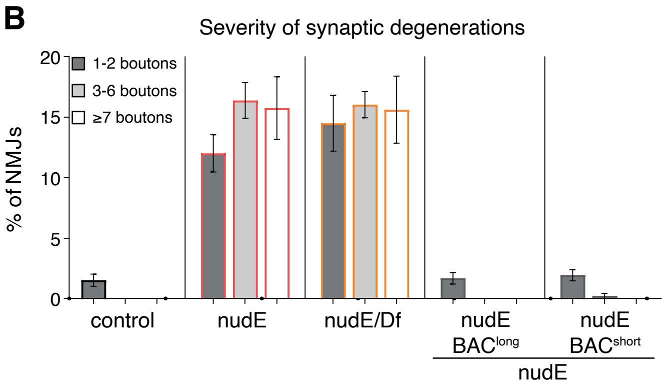


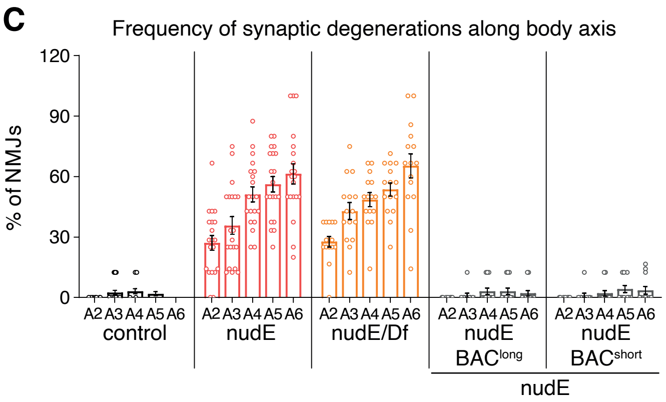


**D**

| Genotype | **Viability** | **Larval movement** |
|---|---|---|
| nudE$^{39A}$ | late larval/early pupal lethal | sluggish to paralyzed |
| nudE$^{39A/Df}$ | late larval/early pupal lethal | sluggish to paralyzed |
| nudE$^{39A}$, BAC long | viable | normal |
| nudE$^{39A}$, BAC short | viable | normal |
| nudE$^{39A}$, da >nudE | viable | normal |
| nudE$^{39A}$, elav >nudE | late larval/early pupal lethal | normal |
| nudE$^{39A}$, ok >nudE | late larval/early pupal lethal | normal |
| nudE$^{39A}$, mef2 >nudE | late larval/early pupal lethal | sluggish to paralyzed |

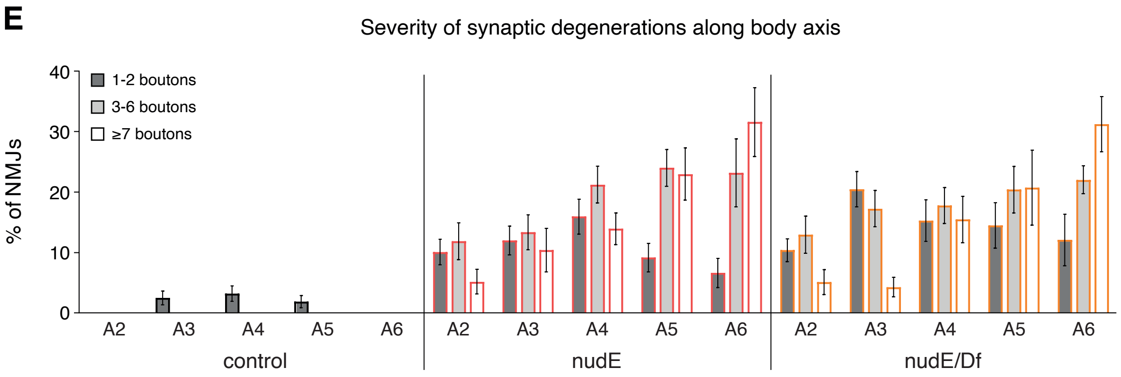

### Supplemental Figure S2

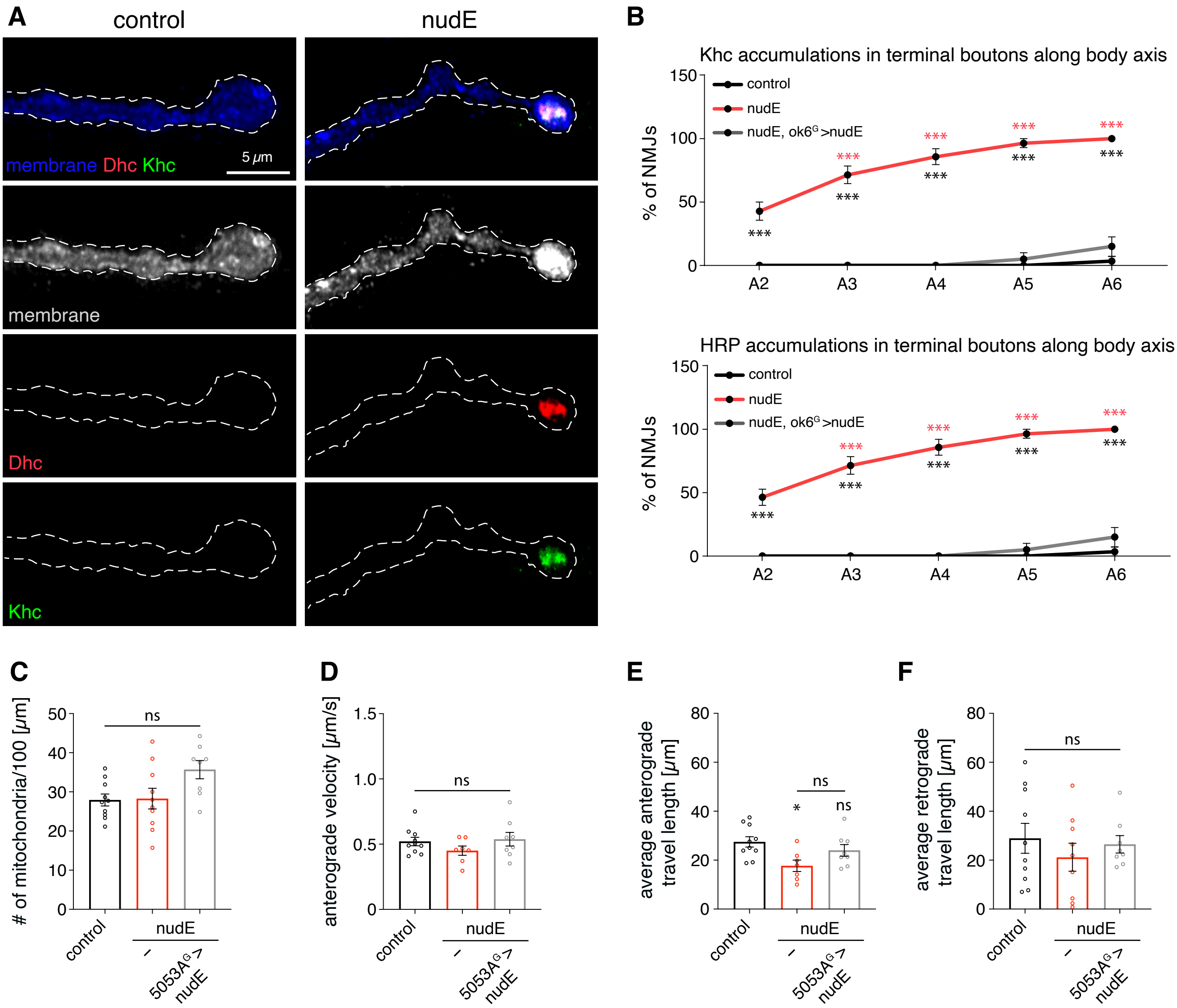

#### Supplemental Figure S3

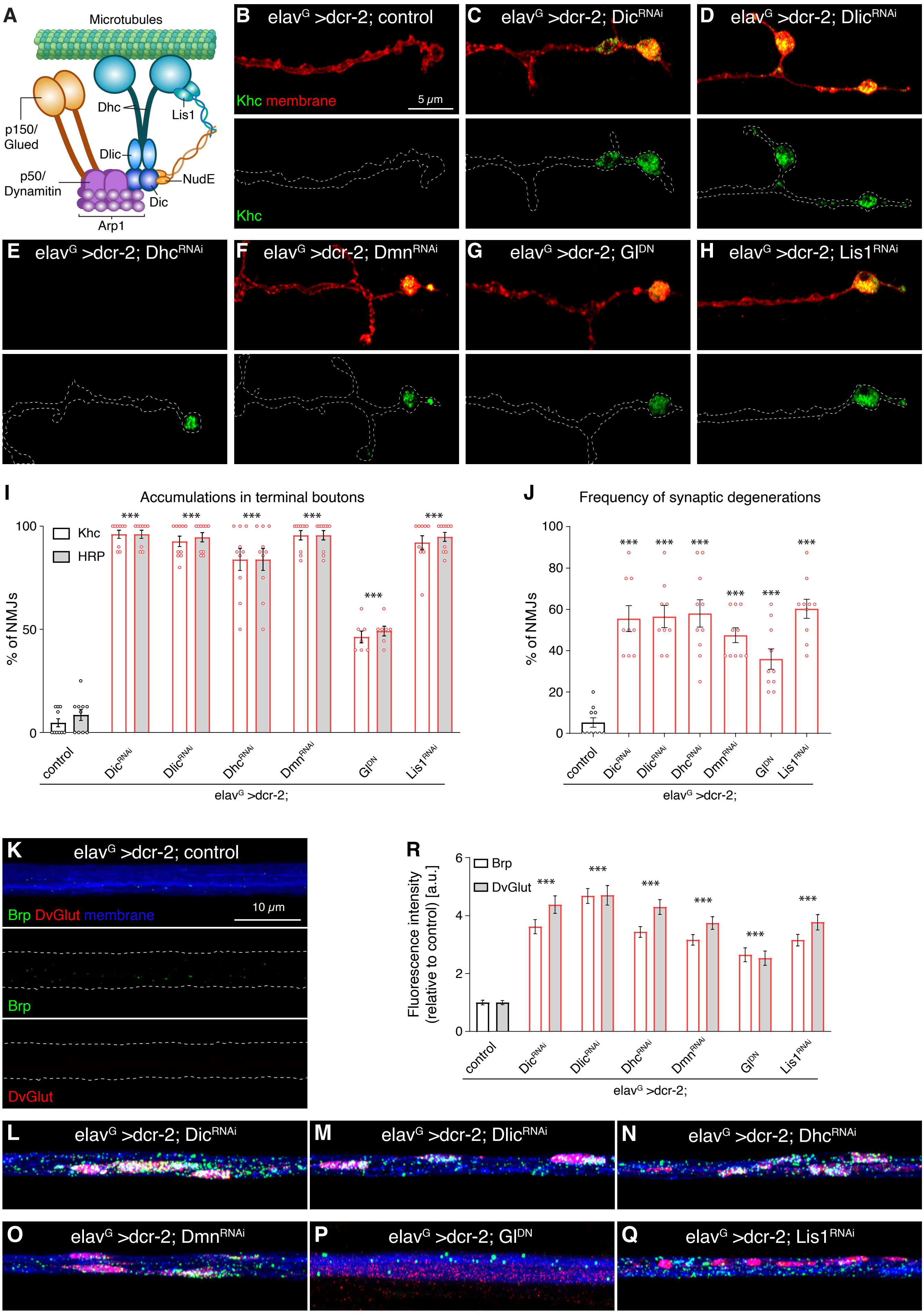

#### Supplemental Figure S4

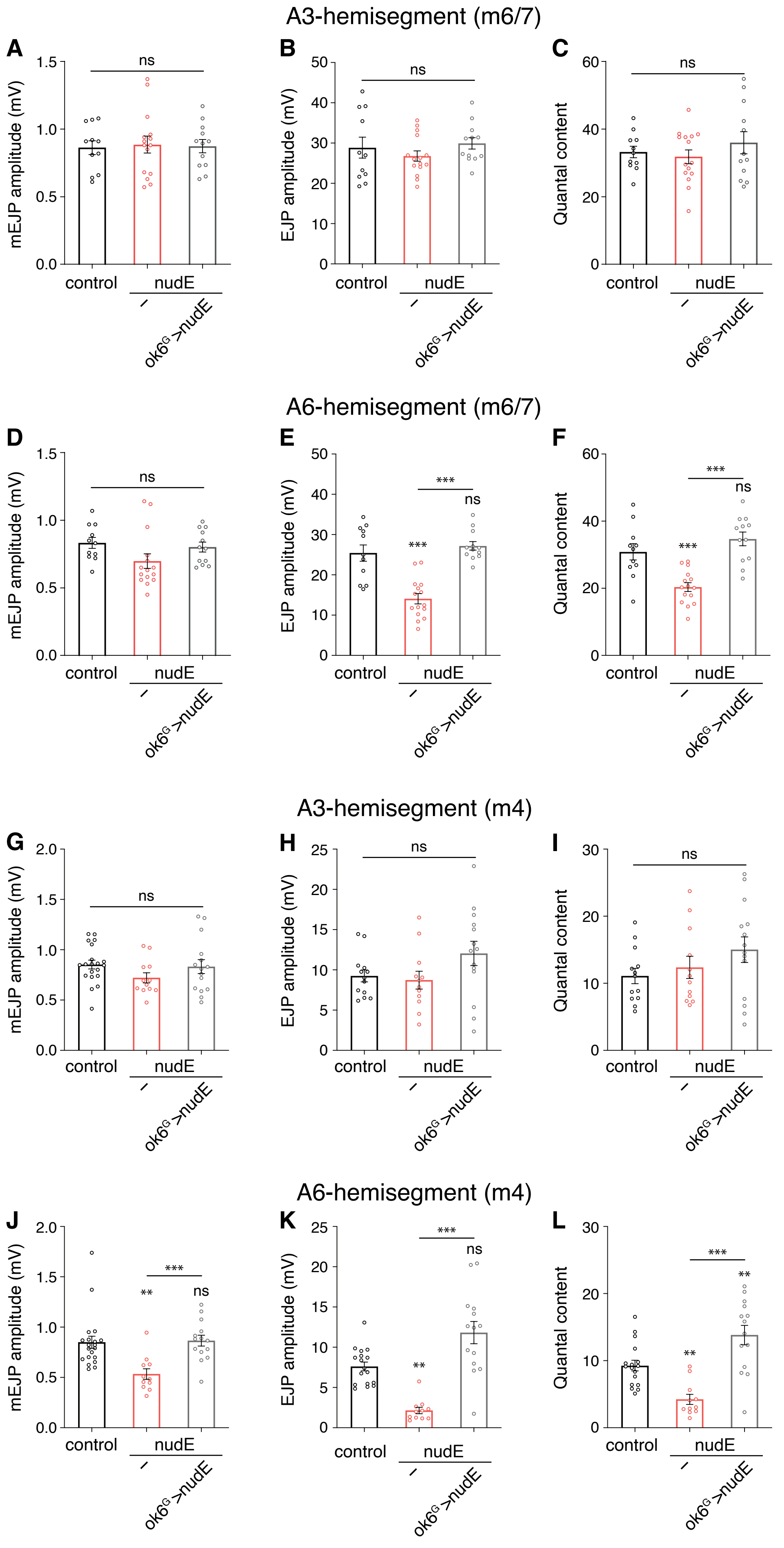

#### Supplemental Figure S5

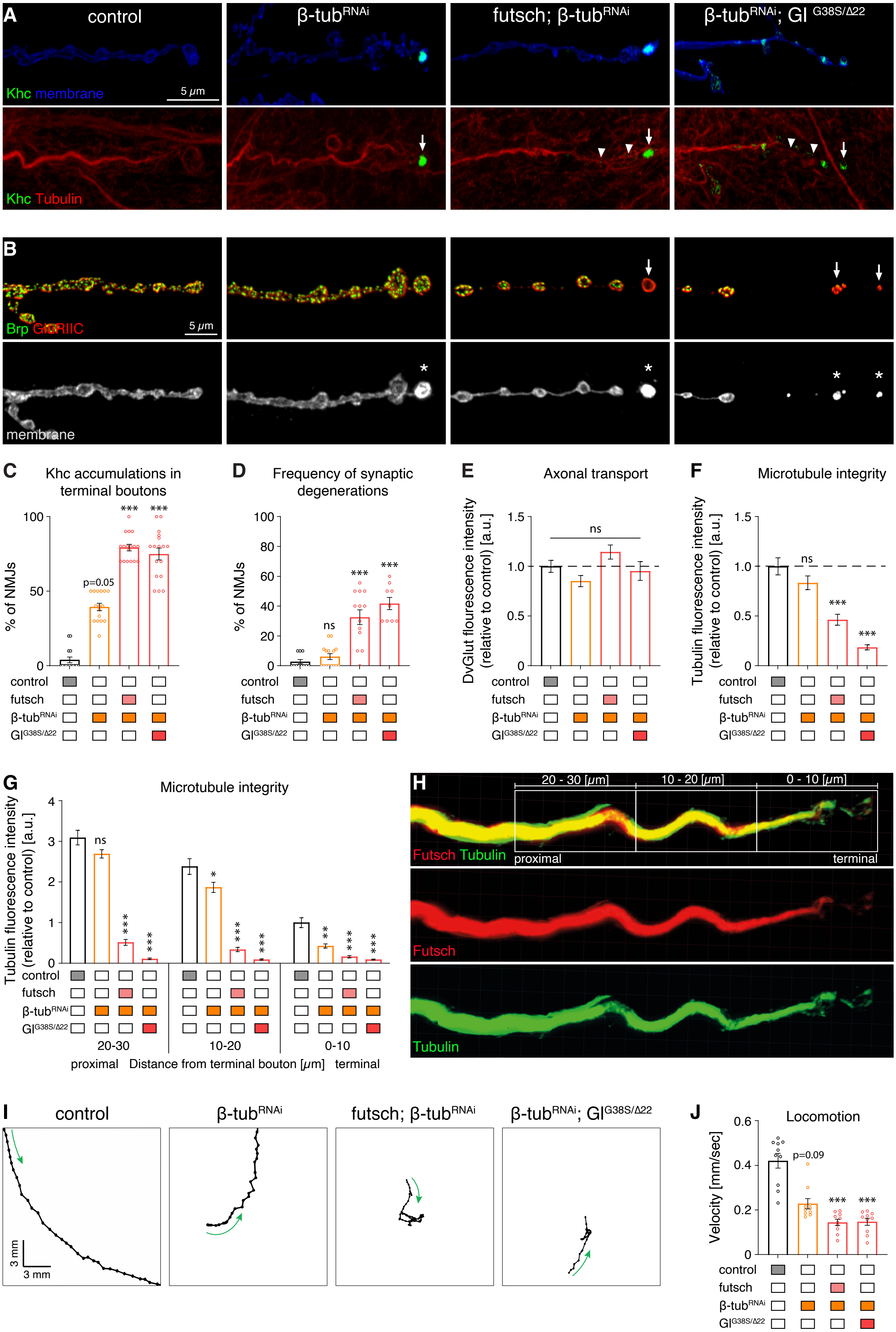
